# A Combined Approach to Extract Rotational Dynamics of Globular Proteins undergoing Liquid-Liquid Phase Separation

**DOI:** 10.1101/2024.09.17.613388

**Authors:** Dominik Gendreizig, Abhishek Kalarikkal, Simon L. Holtbrügge, Saumyak Mukherjee, Laura Galazzo, Svetlana Kucher, Arnulf Rosspeintner, Lars V. Schäfer, Enrica Bordignon

## Abstract

The formation of protein condensates (droplets) via liquid-liquid phase separation (LLPS) is a commonly observed phenomenon in vitro. Changing the environmental properties with cosolutes, molecular crowders, protein partners, temperature, pressure, etc. was shown to favour or disfavour the formation of protein droplets by fine-tuning the water-water, water-protein and protein-protein interactions. Therefore, these environmental properties and their spatiotemporal fine-tuning are likely to be important also in a cellular context at the existing protein expression levels. One of the key physicochemical properties of biomolecules impacted by molecular crowding is diffusion, which determines the viscoelastic behaviour of the condensates. Here we investigate the change in the rotational diffusion of γD-crystallin, undergoing LLPS in vitro in aqueous solutions in absence and presence of cosolutes. We studied its rotational dynamics using molecular dynamics simulations (MD), electron paramagnetic resonance (EPR) spectroscopy and fluorescence spectroscopy. MD simulations performed under dilute and crowded conditions show that the rotational diffusion of crystallin in water is retarded by one to two orders of magnitude in the condensed phase. To obtain the rotational dynamics in the dilute phase we used fluorescence anisotropy and to extract the retardation factor in the condensed phase we used spin-labeled γD-crystallin proteins as EPR viscosity nanoprobes. Aided by a viscosity nanoruler calibrated with solutions at increasing sucrose concentrations, we validate the rotational diffusion retardation predicted by MD simulations. This study underlines the predictive power of MD simulations and showcases the use of a sensitive EPR nanoprobe to extract the viscosity of biomolecular condensates.

## Introduction

The phenomenon of liquid-liquid phase separation (LLPS) evokes a strong interest in the context of biological systems, as a manifold of processes, like the formation of membraneless organelles or the organization of the cytoskeleton, involve the formation of phase separated micro-environments for the full functionality of cellular organisms. ^1–10^ The finely tuned equilibrium between inter-and intramolecular interactions in solution determines the phase separation behavior. The thermodynamic process of phase separation for binary mixtures of solvents and small molecules is well understood but with increasing size and complexity of the molecules it becomes more challenging to disentangle all existing interactions. ^11–13^ Recent studies showed a composition dependency for the propensity to undergo LLPS in polymers and peptides: while charged soluble polymers tend to phase separate with an upper critical solution temperature (UCST), hydrophobic polymers tend to show the opposite behavior by phase separating with a lower critical solution temperature (LCST). ^14, 15^ Furthermore, a manifold of intrinsically disordered proteins (IDPs) follows the same trends and can be approximated by the Flory-Huggins model for charged polymers. ^16^ However, the mechanisms underlying the phase transitions for folded globular proteins are less elucidated. The flexibility of IDPs, small peptides and polymers allows a maximization of intermolecular interactions (protein-protein and protein-water), while the more rigid structures of folded globular proteins constrain these interactions to the solvent-exposed sites. The intra-and intermolecular interactions can be also tuned by cosolutes (e.g. salts, crowders, partner proteins, etc.). ^17–19^ Incorporating cosolutes interacting mostly with the water molecules, such as the osmolyte trimethylamine N-oxide (TMAO), present in the cells of deep-sea organisms, or the crowding agent polyethylene glycol (PEG), was shown to modulate the phase separation behavior of proteins, either folded or unfolded. ^20–23^ This supports the notion that the phase separation propensity is significantly impacted by entropic and enthalpic contributions of the solvent. ^24, 25^

For proteins undergoing LLPS, the coexisting dense and dilute phases have distinct bulk and molecular properties due to their different micro-environments. As recently shown for the IDP system ProTα-Histone 1, macromolecules in the dense phase are exposed to enhanced intermolecular interactions which can increase the bulk viscosity up to 300 times with respect to water, and yet allow extensive sub-microsecond protein chain dynamics in the droplets. ^26^ In the droplets, the change in bulk viscosity, which correlates with changes in translational and rotational diffusion of the proteins, and the regulation of the intra-molecular dynamics are key factors for the physiological function of the condensates in cells. ^27–30^

The protein chosen in this study is the 21 kDa human γD-crystallin (Fig. 1A), the most abundant protein found in the human eye lens (concentrations in the nuclear region > 400 mg/ml). ^31^ It has been structurally characterized in its isolated form (PDB ID: 1HK0) and thoroughly investigated by several biophysical methods during its liquid-liquid phase transition. ^32–35^All members of the γ-crystallin family are soluble globular proteins ubiquitously found among vertebrates and are essential for the maintenance of the refractive index in the eye. ^36, 37^ Due to the nature of their environment in the eye lens with limited protein turnover, the γ-crystallin structures evolved to be particularly stable. ^38^ All γ-crystallins consist of two globular domains with a characteristic Greek key motif containing beta sheets with an unordinary high number of polarizable amino acids leading to a strong difference in refractive index at high concentration and an increased protein density compared to other proteins. ^37–39^

**Figure 1.**
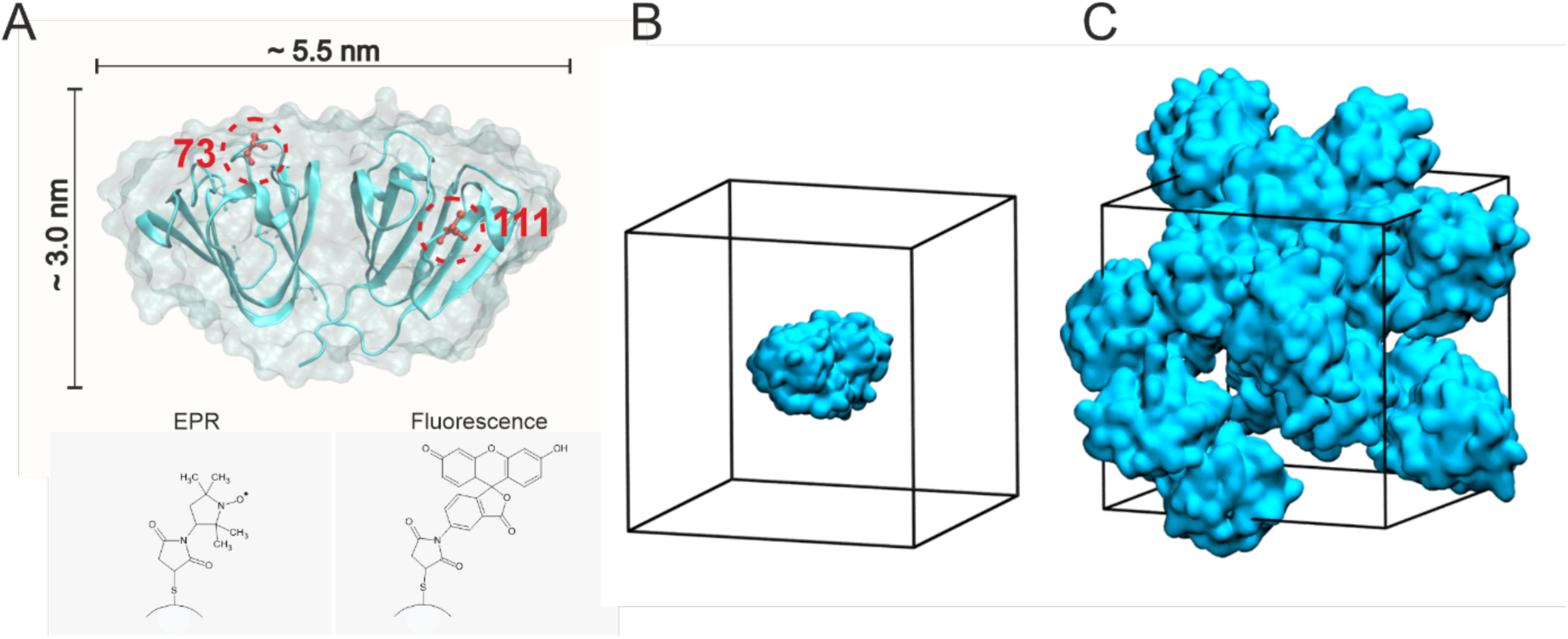
MD simulation system and labeling strategy. (A) X-ray structure of γD-crystallin (PDB: 1HK0) with spatial dimensions indicated. Two positions were selectively labeled for EPR with the maleimido-3-proxyl label (MAP, shown in the inset at the bottom): the native C111 in the wild type and the genetically encoded S73C in the C111S variant, respectively. The native C111 was also labeled with maleimido-5-fluorescein (inset at the bottom) for fluorescence experiments. (B, C) Snapshots of the MD simulation systems used in this work: isolated protein in water (B) and condensate at 420 mg/ml γD-crystallin concentration (C).

Human γD-crystallin presents mainly charged and polar amino acids on its surface, while shielding hydrophobic residues within the Greek key motif in the core of the protein. Intriguingly, misfolding due to a single point mutation induces aggregation of the protein, which has been shown to create a blurring of the eye lens, commonly known as cataract. ^40, 42, 43^ In addition to that, exposition of the eye of fishes and mammals to low temperatures induces the formation of cold cataract, linked to LLPS formation. ^44^

In contrast to IDPs or proteins with intrinsically disordered regions (IDRs) which undergo phase separation already at micro-or even nanomolar concentrations at physiological temperatures, γD-crystallin phase separates at millimolar concentrations. Experimentally it was demonstrated that the UCST for γD-crystallin at 190 mg/ml (∼9 mM) concentration is around 12 -13 °C, which can be tuned by the addition of different cosolutes in a concentration dependent manner. ^33, 34^

One of the major environmental changes that a protein faces while undergoing LLPS is the appearance of molecular crowding, which is expected to affect both its rotational and translational dynamics. Thus, biophysical methods sensitive to changes in the protein dynamics, like NMR, fluorescence spectroscopy and EPR have been used as suitable methods to study these changes. ^20, 26, 45–59^

In this work, we characterize the changes in rotational dynamics of a globular protein γD-crystallin undergoing LLPS in absence and presence of cosolutes using a multidisciplinary approach, including MD simulations, EPR spectroscopy, and fluorescence spectroscopy.

## Computational Methods

### System setup

The initial atomic coordinates of the human γD-crystallin protein were taken from the Protein Data Bank (PDB ID: 1HK0) (Fig. 1). To prepare the dilute system, a single protein molecule was placed in a cubic box with edge length ∼7.6 nm and was solvated with 14014 water molecules (Fig. 1 B). The “condensate” system was prepared by introducing 8 molecules of the protein in a ∼8.7 nm cubic box (Fig. 1). This corresponds to a protein concentration of ∼420 mg mL^-1^, which is the estimated concentration of γD-crystallin condensates at 273 K reported in Mukherjee et al.. ^24^ The 8 proteins were solvated with 15402 water molecules. In both systems, 150 mM sodium chloride (NaCl) was added to mimic experimental conditions. This amounted to 41 Na^+^ and 41 Cl^-^ ions in the dilute system, and 62 Na^+^ and 62 Cl^-^ ions in the condensate system. As the protein does not carry a net charge when all titratable side chains are modelled in their standard protonation states at pH 7, no excess ions needed to be added to neutralize the systems.

### Simulation details

All MD simulations were performed using the GROMACS package (version 2020.6) as previously reported. ^24^ The amber99SB-disp force field was used to describe the protein, water and ions. Virtual sites for H-atoms were used, which enabled integrating the equations of motion with 4 fs time steps with the leap-frog algorithm. ^50^ The energy of both systems was initially minimized using steepest decent. Thereafter they were equilibrated under NpT conditions at constant temperature (273 K) and pressure (1 bar) with harmonic position restraints on the protein heavy atoms. This was followed by an NpT equilibration without any position restraints for a duration of 200 ns. Finally, the systems were simulated under NVT conditions for 10 µs. The last 5 µs of the trajectories were used for the subsequent analyses. Snapshots of the equilibrated systems are shown in Figure 1C.

The temperature and pressure in the simulations were controlled using the Nosé-Hoover thermostat and the Parrinello-Rahman barostat, respectively. ^51, 52^ Non-bonded interactions (Coulomb and van der Waals) were computed up to a distance cut-off of 1.0 nm using a buffered Verlet pair list. ^53, 54^ Long-range electrostatic interactions were treated with the particle mesh Ewald method with a grid-spacing of 0.12 nm. The lengths of the covalent bonds in the proteins were constrained using the LINCS algorithm and SETTLE was used to constrain the internal degrees of freedom of the water molecules. ^55^

### Analyses

To quantify the rotational dynamics, the isotropic rotational diffusion coefficients *D*_iso_ of γD-crystallin molecules were calculated from the MD trajectory following the method of Chen et al. ^56, 57^ First, for each protein in the simulation box the time-dependent orientations *R*(*τ*) were determined in steps of 100 ps along the last 5 µs of the trajectory using the ‘rotmat’ program of GROMACS. The remaining analyses were conducted using our Python package rotationaldiffusion (https://github.com/MolSimGroup/rotationaldiffusion).

The orientations (or rotations) *R*(*τ*) were converted to quaternions, which embody the rotation angle *α* and rotation axis defined by the unit vector 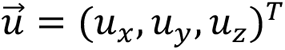
 as

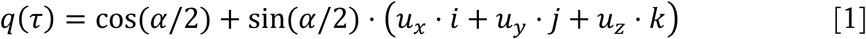

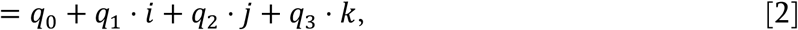

where *i*, *j*, and *k* are distinct imaginary numbers.

Using the inverse (or complex conjugated) quaternions

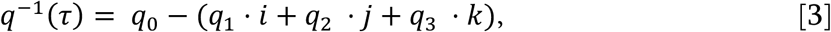

the rotations from time *τ* to time *τ* + *t* were computed as the quaternion products

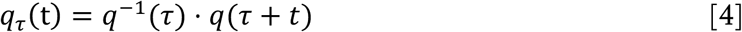

for all possible combinations of *τ* and *t*, up to a maximum lag time of *t* = 500 ns.

The three imaginary parts of rotations *q*_*τ*_(*t*) with rotation angles *α*_*τ*_(*t*) were used to compute the isotropic rotational correlation function

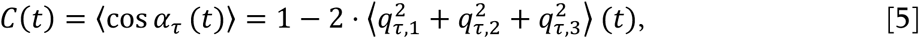

where 〈… 〉 denotes averaging over all starting times *τ*.

*D*_iso_ was finally obtained by fitting the right side of Eq. 5 with the analytical function

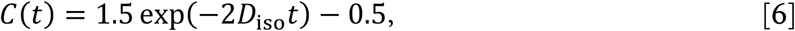

which is a valid model for proteins that behave as rigid rotors undergoing Brownian motion.^52^ This approximation provides accurate rotational diffusion constants for globular proteins in aqueous solution. ^56, 58^

The statistical uncertainty of *D*_iso_ in the dilute system was estimated from the standard deviation of the diffusion coefficients extracted from 100 isotropic Brownian dynamics simulations, which were propagated for 5 µs (similar to the MD simulations). ^58^ Reliable uncertainties could not be estimated for the condensate system due to limited statistics (see Results).

The outlined procedure was used to compute the isotropic diffusion coefficient *D*_iso_ for each of the nine γD-crystallin molecules, one in the dilute system and eight in the condensate system.

## Experimental Methods

### Protein preparation

The wild type protein (molecular weight 21.4 kDa, including the 6x-His tag) was prepared as in Ref. 24. Briefly, a pET-30b(+) vector containing the CRYGD gene for human γD-crystallin with a C-terminal 6x-His tag was transformed into BL21DE3-RIL cells. The cells were then grown at 37°C until OD_600_ of 2.0 was reached and later induced with 0.5 mM IPTG and grown overnight. The cells were then harvested by centrifuging at 5000 rpm for 15 min in the Thermo scientific Sorvall Lynx 6000 centrifuge. For purification of wild type γD-crystallin, the cells were lysed through sonication in Lysis buffer (50 mM NaH2PO4 + 750 mM NaCl + 2.5 mM Imidazole + 2 mM TCEP, pH = 8.0). The His-tagged protein was purified by Affinity chromatography using a manually packed column containing HisPur Cobalt Resin (Thermo Scientific). Size exclusion chromatography was performed with the HiLoad 26/60 Superdex 75 with 20 mM TRIS + 150 mM NaCl pH=7.5 with 2 mM TCEP as the elution buffer buffer. The protein was concentrated using the Vivaspin 20 columns, 10000 MWCO (Sartorius). The concentration was measured by absorbance at 280 nm (ε = 42860 M^-1^cm^-1^ for the protein with the 6x-His tag). Aliquots of the protein solution were stored at -20 °C after flash-freezing in liquid nitrogen.

The site-directed mutation was carried out through PCR. Initially, the C111S mutation was created using the 5’ TTT ACC GAA GAC TGC AGC TCC CTG CAA GAT CGT TTC 3’ forward and the 5’ GAA ACG ATC TTG CAG GGA GCT GCA GTC TTC GGT AAA 3’ reverse primer. Then the plasmid with the C111S mutation was used as the template for inserting the S73C mutation using the primers 5’ GAT GGG CCT GTC CGA CAG CGT TC 3’ forward and 5’ GAA CGC TGT CGC ACA GGC CCA TC 3’ reverse primer. The C111S-S73C mutant was purified as the wild type.

### Spin labeling

γD-crystallin wild type (C111) and the mutants S73C and C111S-S73C were incubated with 1 mM DTT for 1 h at room temperature. DTT was removed by a PD-10 desalting column. 3-Maleimido-2,2,5,5-tetramethyl-1-pyrrolidinyloxy (MAP) (Sigma Aldrich) was added to the single cysteine variants (C111 and C111S-S73C) in a 1:1.2 ratio, while the double mutant (S73C) was incubated with MAP in a 1:2.2 ratio. The samples were incubated overnight at 4°C and in the dark under constant mixing. Excess label was removed via PD-10 desalting column. The proteins were concentrated via Vivaspin 20 column with a 10 kDa cutoff and the final concentration was determined via absorbance at 280 nm (ε = 42860 M^-1^cm^-1^). The labeling efficiency was determined via continuous wave EPR against a standard of 100 µM TEMPOL in water by double integration using the free software spintoolbox (https://www.spintoolbox.com/en/). The labeling efficiencies were determined to be 120±10 % spin/protein for the mutant C111-S73C, 100±10 % spin/protein for the mutant C111S-S73C and 70±10 % spin/protein for the wild type C111.

### Continuous wave (cw) EPR

All cw EPR experiments were performed on a Bruker E500 X-band equipped with a super high Q cavity ER 4122 SHQ. The microwave power was set to 2 mW, the modulation amplitude to 0.1 mT and the temperature was controlled with a nitrogen flow cryostat and a temperature controller (Eurotherm). The series of spectra at different concentration of sucrose (w/v) were detected on the spin-labeled crystallin C111S-S73C or C111 (50 μM) at 293K.

The samples used to determine the onset temperature of LLPS were prepared by adding 50 µM spin-labeled protein of the respective mutant to 2 mM unlabeled γD-crystallin and the respective cosolute in a total volume of 20 µL to 50 µL glass capillary (Blaubrand). Samples containing 50 µM spin-labeled protein with cosolutes were used as reference. The temperature was set with a in a range from 5 °C to 35 °C in steps of 5 °C. The spectra were acquired from 5 °C to 35 °C and the reversibility of the spectral changes was confirmed after the temperature sweep.The spectral changes of the temperature-dependent cw EPR spectra were evaluated with MATLAB 2022. All spectra were spin normalized to the second integral (https://www.spintoolbox.com/en/). The difference spectra were created by subtraction of the spectrum in the presence of cosolute and 2 mM unlabeled protein and the spectrum of the reference sample containing the same cosolute. All spectra were splined before subtraction. To obtain the onset temperature of LLPS from the difference spectra, the absolute integral of the low-field region was plotted against the temperature.

To identify the spectral features of the condensed phase, we incubated a large sample volume 60 µl in a glass capillary containing the spin-labeled crystallin C111S-S73C or C111 (50 μM) in the presence of 2 mM unlabeled protein and 0.6 M TMAO at 293 K (LLPS conditions). We gently centrifuged the sample for ca. 20 seconds using a hand centrifuge, thereby creating a clear separation between the dilute phase on the top and the condensed phase at the bottom of the tube. The spatial separation of the two phases in the tube allowed us to fill the EPR cavity with the dilute phase (upper part of the tube) and with most of the condensed phase (bottom part of the tube). The spectrum of the dilute phase (upper part) was compared with the series of spectra obtained with different sucrose content on the same mutant to obtain the viscosity that best corresponded to the dilute phase conditions. To obtain the spectrum of the pure dense phase we proceeded as follows: we subtracted from the spectrum of the dense phase (bottom part of the tube) increasing fractions of the spectrum of the dilute phase (upper part of the tube). The normalized difference spectrum was compared with a normalized spectrum of known viscosity (different percentages of sucrose) and the residual was calculated. By plotting the absolute integral of the residual versus the fraction of the dilute phase removed for each sucrose content analyzed, we obtained the optimal fraction of the dilute spectrum and the viscosity which best represented the pure dense phase (Fig. 5 and Fig. S8).

### Pulse EPR

The pulse EPR experiments were performed on the C111, S73C-C111S and S73C-C111 spin-labeled proteins with a Bruker Q-band Elexsys E580 spectrometer equipped with a 150 W TWT amplifier and a Bruker Spinjet-AWG at 50K. A homemade probe head accommodating 3 mm o.d. tubes was used. ^59^

For Gaussian pulses, the predefined pulse shape (Shape 1) in Bruker Xepr version 2.6b.170 was used. As standard setup we used the dead-time free 4-pulse DEER sequence with 16-step phase cycling. ^60, 61^ All pulses were set up to the same length (32-34 ns corresponding to 16-20 ns FWHM) The pump frequency was positioned at the maximum of the nitroxide spectrum and the observer at –90 or -100 MHz depending on the DIP of the resonator. Table 1 summarizes the additionally used parameters.

**Table 1:** Applied DEER parameters.

|  |  |
| --- | --- |
| Shot repetition time [ms] | 4 |
| $t_1$ [ns] | 400 |
| $t_2$ [ns] | 3000-5000 |
| Zero time [ns] | 120 |
| $t_1$ averages in steps of 16 ns | 8 |

Three DEER samples (40µL, 3 mm o.d. quartz tubes from Aachener Quarzglas) were prepared: two control samples at micromolar protein concentration where no LLPS is present before flash freezing and one sample at 2 mM protein concentration enriched in condensates. Sample description: 1) 50 µM of the S73C-C111 spin-labeled protein with 50% deuterated glycerol (Sigma Aldrich), sample shock-frozen from room temperature in liquid nitrogen, no turbidity observed before freezing; 2) 50 µM of the S73C-C111 spin-labeled protein with 5% of PEG6000, sample shock-frozen from room temperature in liquid nitrogen, no turbidity observed before freezing; 3) 50 µM of the S73C-C111 spin-labeled protein mixed with 2 mM of unlabeled protein and 5% of PEG6000. The sample was incubated at 4°C (the presence of condensate was evident by the turbidity) for 30 minutes prior to shock-freezing in liquid nitrogen. In the second experiment, to minimize the signal arising from the dilute phase, the tube was centrifuged using a hand centrifuge (Hettich AG) after the incubation, the clear solution was quickly removed using a Hamilton tubing on ice, and the dense phase was immediately frozen. The DEER data were analyzed with DeerAnalysis 2021. ^62^

### Labeling of γD-crystallin with fluorophores

Fluorescein-5 Maleimide (62245) was purchased from Thermo Scientific. 100 μM of γD-crystallin wild type was first reacted with 1 mM DTT for one hour. The excess DTT was removed by passing the protein solution through Econo-Pac 10G desalting column (Biorad). The concentration of the protein solution was measured via UV-vis absorption at 280 nm and the fluorophore was added in a 1:1 protein-fluorophore molar ratio for labeling. The protein solution was kept stirring in the cold room (at 6 °C) for 16 hours. The solution was passed through a PD-10 column to remove any unbound fluorophore. The labeled protein was concentrated to the desired concentration using Vivaspin Turbo 4 10000 MWCO concentrators (Sartorius). The protein concentration was determined with the formula

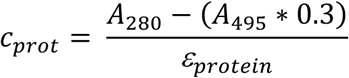

to account for the absorption of fluorescein at 280 nm (https://www.aatbio.com/resources/correction-factor/fluorescein).

The labeling efficiency was subsequently determined by

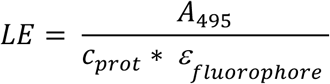

### Fluorescence lifetime and time-resolved anisotropy

The fluorescence lifetimes and anisotropy decays were measured at room temperature using a home-built set-up described in reference. ^63^ In particular, a pulsed laser diode at 437 nm (PicoQuant, LD-440) was used as excitation source and a cooled photomultiplier (PicoQuant, PMA-C-192-N-M) as detector.

For all samples 520 nm was used as detection wavelength. The measurement time was the same for all measurements. Magic angle measurements were performed at the start and the end of each sample measurement series to check for sample decomposition/changes, which were not observed. To determine the g-factor of the detection set-up, a sample of fluorescein in water was measured and tail-matching to the parallel and perpendicular data applied

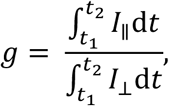

where t_1_ is the time where the anisotropy has fully decayed (here we chose 3 ns) and t_2_ is chosen to limit the data to a range with relevant photon counts (10 ns).

The anisotropy, *r*, was calculated according to

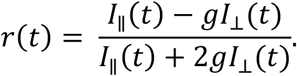

As we are merely interested in the long-time anisotropy decay we decided to fit a multiexponential model to the anisotropy data at times longer than 0.5 ns, not accounting for the finite duration of the exciting laser-pulse. We also limited the fitting window to times shorter than 20 ns, because a reflection in the fiber renders the data, showing up at longer times, useless. We performed a weighted nonlinear least-squares fit optimizing the reduced chi-square, χ^2^_r_, as defined via

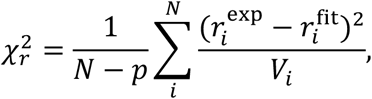

where *p* is the number of fitting parameters, *N* the total number of data points, and where the variance of the anisotropy, *V_i_*, is obtained by error propagation and is given by^64^

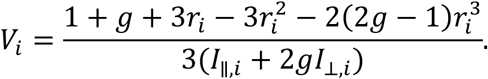

Time zero of the time-dependence of the anisotropy was found by taking the maximum of the instrument response function and accounting for the detector induced color-shift.

## Results and Discussion

### Rotational diffusion of the proteins in dilute and condensed phase from MD simulations

The isotropic diffusion coefficients *D*_iso_ and rotational correlation times *τ*_c_ = 1/6*D*_iso_ of γD-crystallin at 273 K were determined using MD simulation trajectories of dilute and condensate systems from our prior study. ^53^ For the present work, the isotropic rotational correlation functions of the γD-crystallin molecules were extracted from the simulations (Fig. 2a, dashed lines) and fitted by using Equation 6 (Fig. 2a, solid lines). For the dilute system, this yielded a rotational correlation time of 30 ± 3 ns.

**Figure 2.**
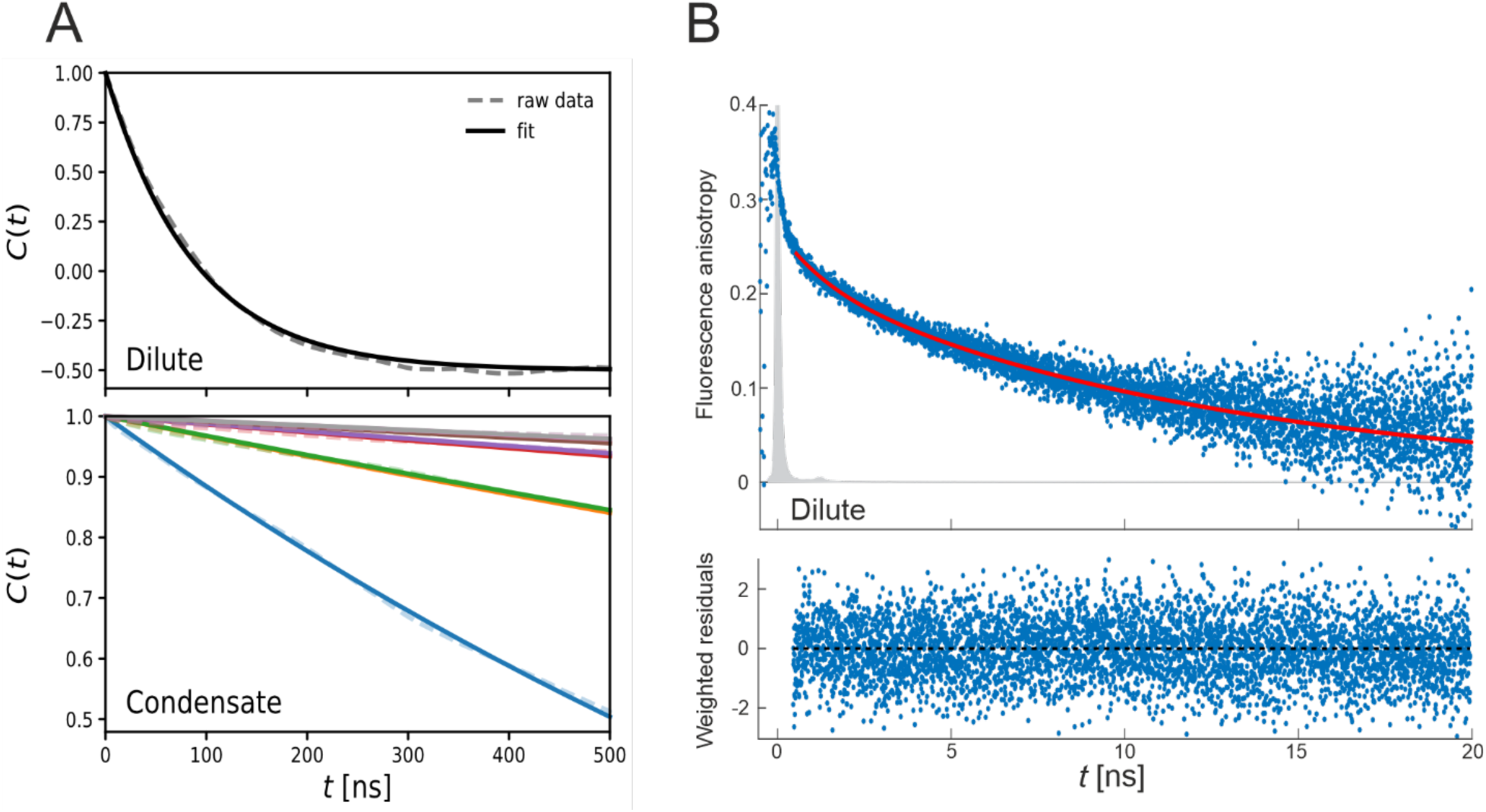
Rotational dynamics from MD simulations and fluorescence anisotropy. A) Rotational correlation functions of the proteins in dilute (top) and condensate systems (bottom) from MD simulations at 273 K. The dotted lines represent the raw data and the solid lines denote exponential fits. In condensates the correlation functions and fits for all the 8 proteins in the system are shown. B) Experimental fluorescence anisotropy measurement of 200 nM fluorescein-labeled γD-crystallin in the dilute phase at 293 K. The anisotropy function consists of two components. To extract the rotational correlation time only the long component was fitted (solid line) with a monoexponential function resulting in rotational correlation time of 12.2 ± 0.2 ns.

The condensate system contains eight γD-crystallin molecules in the simulation box, corresponding to a protein concentration of 420 mg/mL. ^23^ The rotational correlation time *τ*_c_ was determined separately for each protein molecule (Tab. S1), ranging from 6580 ns (grey curve in Fig. 2a) to 415 ns (blue curve in Fig. 2a). Thus, the main finding from the MD simulations is that the rotational dynamics of γD-crystallin in the condensate are strongly retarded compared to dilute conditions, due to proteins hindering each other’s movement. The median correlation time of 3850 ns in the condensate is more than two orders of magnitude slower than the 30 ns found in the dilute phase. Even the least retarded individual protein molecule (*τ*_c_ = 415 ns, see above) is already slowed down by more than one order of magnitude, which we interpret as a lower bound for the retardation. This finding is in qualitative agreement with previous reports, for example, the strong slowdown of tumbling times with increasing protein self-crowding measured by NMR relaxation and fluorescence correlation spectroscopy. ^24^

In addition, rotational correlation times of the eight individual proteins in the condensate simulation system are highly heterogeneous, spanning more than one order of magnitude. This is due to the distinct local environment that each protein experiences during the multi-microsecond MD simulation, i.e., on the timescale relevant for rotational diffusion. Proteins that are in a locally more crowded region, and thus experience stronger interactions with neighboring proteins, are slowed down more strongly (Fig. S1). In the limit of an infinitely long MD trajectory, the rotational correlation times of all individual proteins in the condensate system will be identical, since the proteins are identical. In this sense, capturing the heterogeneous broadening of rotational correlation times due to spatial differences in local protein concentration on short timescales is a feature of a non-converged trajectory and cannot be directly related to spatial inhomogeneity of multiphasic condensates.^65^ This limited sampling also prohibits estimating meaningful statistical uncertainties for the rotational correlation times in the condensate, but the spread of the individual *τ*_c_ inform about the statistical precision. However, an advantage of the MD simulations is that they enable one to probe the individual protein molecules in a (transiently) different local microenvironment, which might also be present in vitro or in the biological cell.

### Rotational dynamics of the dilute proteins by fluorescence anisotropy

To validate the rotational correlation times extracted from our MD simulations, we performed time-resolved anisotropy of crystallin in the dilute phase (200 nM) in aqueous buffer at 293 K. The data analysis unveiled two components, characterized by short and long rotational correlation times. A simple biexponential function provided a rotational correlation time for the slow component, assigned to the crystallin in dilute phase, of τ_c_ = 12.2 ± 0.2 ns (Fig. 2B). The obtained value agrees with the approximation for a globular protein of about 21 kDa in water at room temperature when considering an average density of γD-crystallin of 1.4 g/cm^3^ and assuming a simple Brownian motion. ^35, 64, 65^ Under this approximation, the τ_c_ in ns of a monomeric protein in a room-temperature aqueous solution is approximately 0.6 times its molecular weight in kDa (21.4 kDa for crystallin would correspond to about 13 ns). To compare the experimental value at 293 K (12.2 ns) with that obtained from the MD simulations at 273 K (30 ns), we need to consider that the dynamic viscosity of water at 273 K is 1.785 times higher than at room temperature (and hence the MD-calculated rotational diffusion extrapolated at 293 K would be ∼16.8 ns), which is in the experimentally determined range. ^64^ Therefore, we conclude that the *in silico* analysis accurately captures the rotational dynamics of the protein in the dilute phase.

### Effects on the protein rotational dynamics by EPR

The large retardation for the overall tumbling of the proteins predicted by MD simulations could not be addressed by fluorescence anisotropy in our system due to several limitations such as the multiexponential character of the anisotropy traces, the too fast lifetime of fluorescein (Fig. S2), and the heterogeneous composition of the sample (dilute and condensed phases in equilibrium). Nitroxide probes are sensitive reporters of rotational correlation times in a three-order of magnitude time window, from ca. 30 ps to ca. 100 nanoseconds, and are sensitive to temperature and viscosity changes in their microenvironment (Easyspin simulations and effects of temperature and viscosity on the spin label MAP are shown in Fig. S3). ^66^ Therefore, we attempted to use EPR to detect the changes in the rotational dynamics of micromolar concentrations of spin-labeled crystallin proteins used as viscosity reporters in samples containing millimolar wild type crystallin proteins transitioning from the dilute to the condensed phase. The rotational dynamics encoded in the continuous wave EPR spectra of nitroxide-labeled protein are a convolution of the rotational dynamics of the spin-labeled side chains, backbone motions, and overall tumbling of the protein. ^67–69^ Thus, we expect to have distinguishable spectral shapes for the protein in the two distinct environments, which could be related to the self-crowding.

The proteins undergo LLPS below 5 °C in water at a concentration of 2 mM (Fig. S4), therefore, we decided to use as reference the EPR spectra detected from 5 to 35°C (no LLPS) in samples containing 2 mM of proteins. First, to identify spectral changes due to LLPS, we increased the overall protein concentration (to 5.7 mM) to increase the onset temperature to 13 °C, but the crowding effects on the spectral shapes due to the high millimolar concentrations masked spectral features possibly associated with droplet formation (Fig. S5). Hence, we decided to add cosolutes (PEG and TMAO) to the 2 mM protein sample, which are known to stabilize droplets above 15°C (Fig. S4) and compare the spectra of the spin-labeled proteins in the presence of cosolutes with those of the 2mM reference spectra without cosolutes. The comparison with the reference spectra highlighted the appearance of a characteristic feature at the LLPS onset temperature determined by turbidity assay (Fig. S4).

The emerging spectral features are characterized by a positive peak in the low field region (Fig. 3 and Fig. S5) which indicates the appearance of a nitroxide population with slow motion. The appearance of such population was found to be correlated with the LLPS onset temperatures in presence of different cosolutes (dotted vertical lines in Fig. 3 correspond to the onset temperature determined by turbidity assay shown Fig. S4), confirming that the observed spectral changes are related to the dynamics of the proteins in the condensed phase.

**Figure 3.**
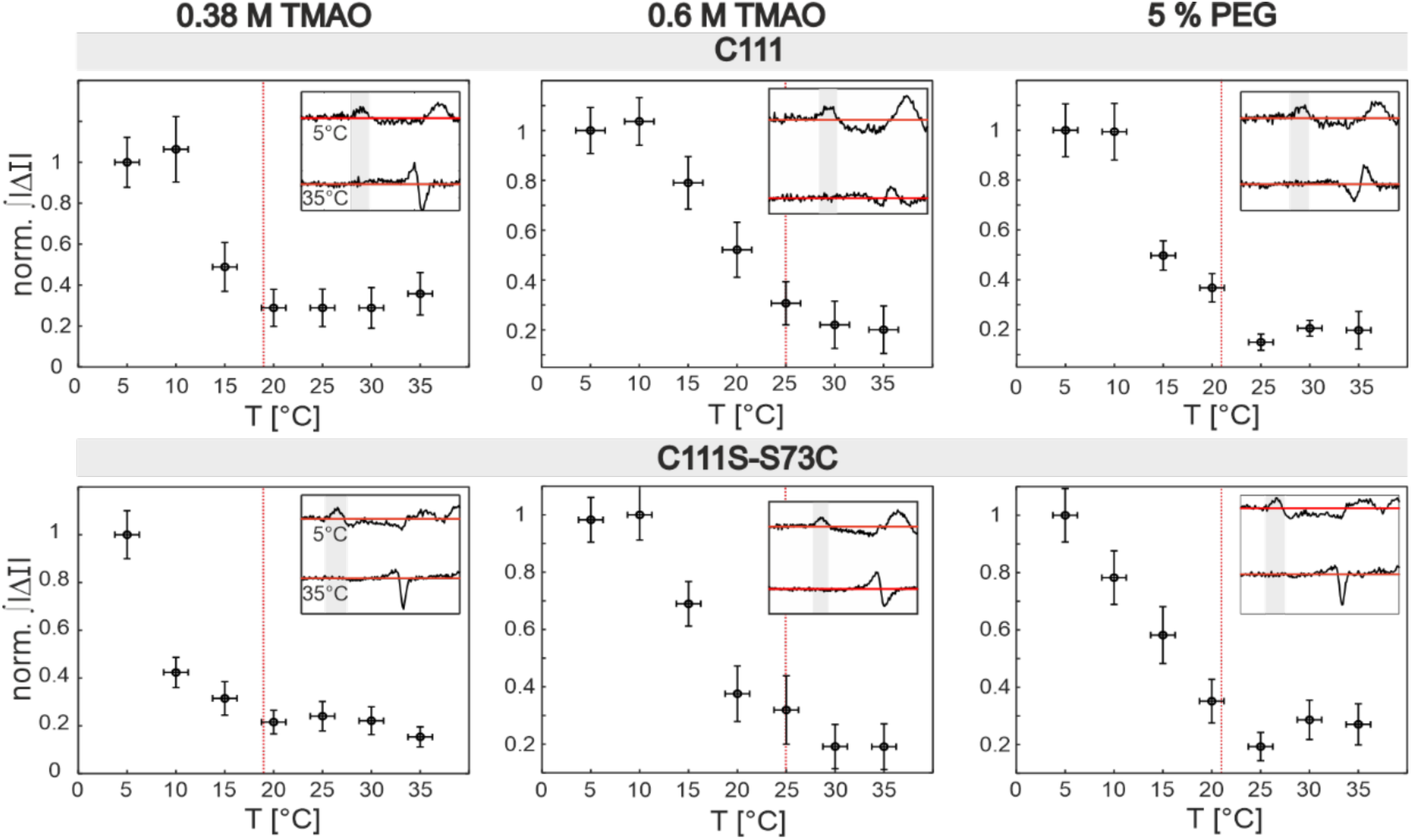
LLPS onset temperature by EPR. The panels show the normalized absolute integral of low field spectral region (gray areas in insets) of the difference spectra of samples containing 2 mM protein in the presence of cosolute minus control as a function of temperature. The samples contain 2 mM wild type γD-crystallin and 50 µM spin-labeled γD-crystallin (top: position C111; bottom: position S73C-C111S) with 0.38 M TMAO, 0.6 M TMAO and 5 % (w/v) PEG as cosolutes. Residual diversion from zero at 35 °C originate from differences in viscosity induced by the cosolute. The inset in each panel compares the difference spectra in the low field region at 5°C and 35°C, the solid red line shows the zero crossing. The red dashed vertical line represents the onset temperature of LLPS determined by turbidity assay (Fig. S4) performed on samples containing 2 mM wild type γD-crystallin with the same solvent composition.

Notably, all cw EPR spectra detected at temperatures below the onset T contain two distinct contributions (condensed and dilute phases), each being a convolution of the global rotation of the protein, the dynamics of the protein segment to which the label is attached, and the motion of the label itself. To further characterize the spectral features of the proteins in the droplets, we physically separated the dilute from the condensed phase via centrifugation directly in the EPR tube (see Materials and Methods) containing 0.6 M TMAO at 20°C. The two resulting spectra are shown in Figure 4A. The spectrum in the dilute phase presents two spectral features, indicative of anisotropic rotational dynamics at the chosen site. An accurate simulation of the spectra would require a multifrequency analysis to infer the exact anisotropic potential at the spin-labeled site under the two different conditions, which is beyond the scope of this work. However, the spectral shape of the dilute phase can be approximated by two isotropic components with rotational correlation times of 2.5 ns (33%) and 12 ns (67%) (black in Fig. 4B, Table 2). The spectrum in the dense phase (Fig. 4A, black) shows distinct features characteristic of a nitroxide approaching the rigid limit of motion (arrows, see Easyspin simulations in Fig. S3A), with the positive peak at the low field region in the condensed phase being the fingerprint feature that we previously identified from the difference spectra (see Fig. 4, Fig. S5 and Table 2). Additionally, we observed a splitting of the negative central peak, indicative of rotational dynamics approaching the rigid limit (see Easyspin simulations in Fig. S3A).

**Figure 4.**
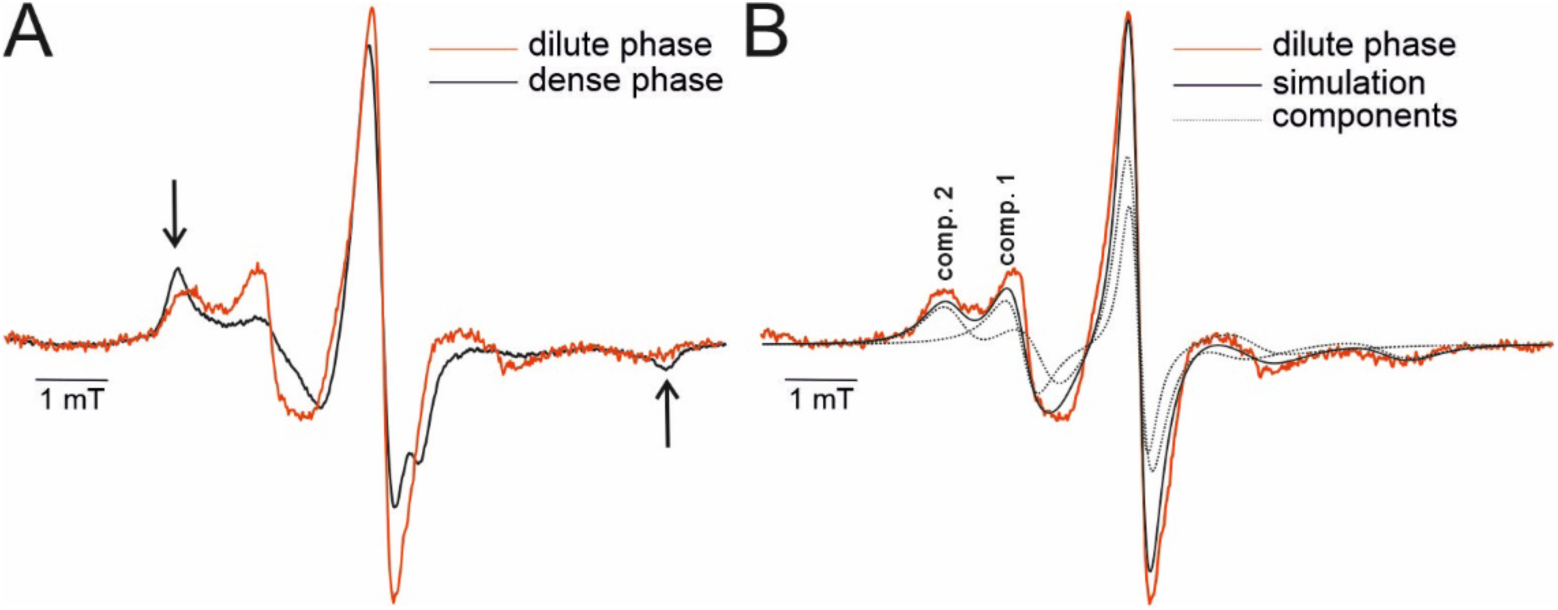
Spectral fingerprint of the dilute and dense phases in equilibrium. A) Cw X-band EPR spectra of γD-crystallin (C111S-S73C) at 293 K recorded on the sample after centrifugation (50 μM spin-labeled protein, 2 mM total protein concentration, 0.6 M TMAO, see Materials and Methods). In red the spectrum of the dilute phase detected on the upper part of the tube, in black the spectrum detected on the bottom part of the tube, containing the dense phase as major component. Arrows highlight the spectral features characteristic of the rigid limit. B) Cw EPR spectrum of γD-crystallin in the dilute phase (red) at 1.5 mM concentration compared to the Easyspin simulations (chili, black) using two isotropic components (dotted black) with rotational correlation times of 2.5 ns (component 1, 33% of the area) and 12 ns (component 2, 67% of the area) (simulation details are shown in Table S3)

**Table 2.** Rotational correlation times and viscosities in the two phases.

| phase | MD<br>at 273 K<br>(extrap. at 293) | Fluorescence<br>anisotropy at<br>293 K | EPR Easyspin at<br>293 K | EPR viscosity<br>nanoprobe at 293 K |
| --- | --- | --- | --- | --- |
| dilute | $\tau_c = 30$ ns (one isolated<br>protein)<br>**17 ns at 293 K** | $\tau_c = 12.2$ ns<br>(at 200 nM) | 12 and 2.5 ns<br>(at 1.5 mM) | $\eta = 1.00$ mPa·s<br>(at 50 μM)<br>$\eta = 1.59$ mPa·s<br>(at 1.5 mM) |
| condensed | $\tau_c = 3850$ ns<br>(415 - 6580 ns)<br>$\tau_c = \mathbf{2200}$ ns at 293 K<br>(240 - 3800 ns) | n.d. | Rigid limit | $\eta > 15.9$ mPa·s |
| Viscosity ratio<br>droplet to dilute<br>($\leq 50$ μM) | >15...220 | n.d. | n.d. | >15 |

n.d.: not determined.

The rotational correlation times of the isotropic components used to simulate the spectra are only indicative but difficult to interpret due to the real anisotropic nature of the labels’ dynamics, and the spectra of the spin-labeled crystallin in the condensate are approaching the rigid limit, thereby impeding the extraction of reliable rotational correlation times. Therefore, to gain insights in the change of viscosity transitioning from the dilute to the condensed phase, we decided to use the spin-labeled crystallin protein as a viscosity ‘ruler’ due to the highly sensitive response of the spectral shape in microenvironments with different viscosities. We measured a series of cw EPR spectra of a spin-labeled crystallin mutant in solutions of known viscosity (aqueous buffer in presence of increasing amount of sucrose, Figure 5A). In this way, the anisotropic spectral features of the ‘viscosity nanoprobe’ can be used as templates to infer the overall viscosity in the dilute and condensed phases. Interestingly, the spectrum of the spin-labeled crystallin in the dilute phase overlaps perfectly with that of a 20% w/v sucrose solution (1.59 higher viscosity than water) (Fig. 5B and Table 2). We confirmed that the increased viscosity with respect to pure water is due to the high protein concentration of 1.5 mM (33 mg/ml) in equilibrium with the droplets (determined by UV-vis absorption). Indeed, molecular crowding effects are already visible in the spectra in the absence of LLPS for protein concentrations > 1 mM (Fig. S7).

**Figure 5.**
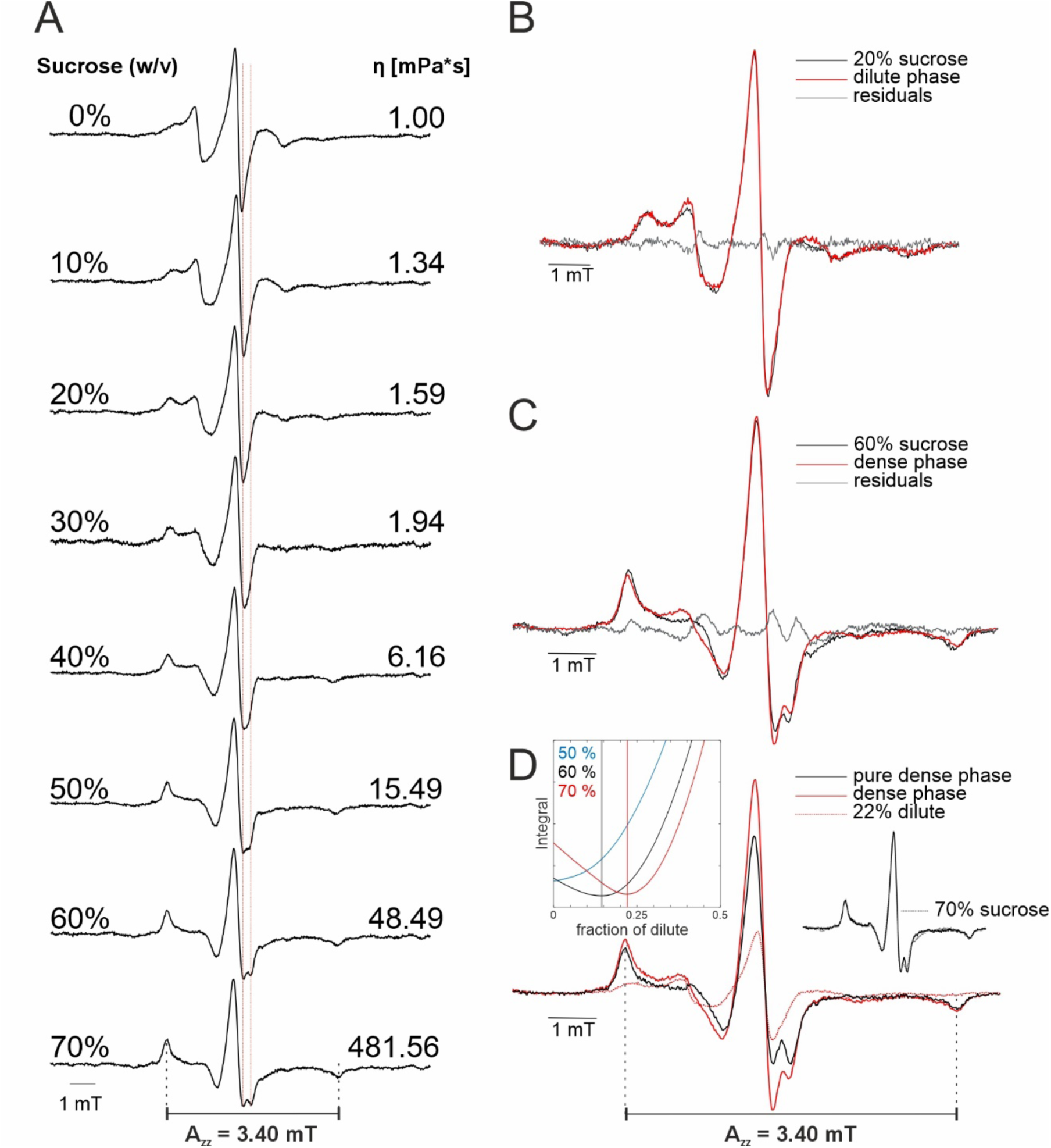
Viscosity ‘ruler’. A) Cw EPR spectra of γD-crystallin (C111S-S73C) recorded at 293 K in aqueous buffer with increasing sucrose concentrations. The corresponding tabulated dynamic viscosity of sucrose solutions at 293 K is indicated on the right. The dashed red line visualizes the splitting of the negative central peak for motions approaching the rigid limit. The value of A_zz_ (reporting on the polarity of the nitroxide microenvironment) was extracted from the spectrum with the highest viscosity to be 3.40 mT. B,C) cw EPR spectra of γD-crystallin in the dilute and dense phase in equilibrium (red) compared to the spectrum of the same mutant in sucrose solutions (black, from panel A) and the residual (grey). The best correspondance for the spin-labeled protein in the dilute phase (concentration determined by UV/vis to be 33±1 mg/ml, 1.5 mM) is found with the spectrum corresponding to a dynamic viscosity of 1.59 mPas. The optimal correspondance for the dense phase is found with the spectrum corresponding to 60% w/v sucrose. D) From the dense-phase spectrum (red, same as in panel C) 22% of the dilute phase spectrum was subtracted (red dotted, as in panel B). The obtained difference spectrum (black) yields the so-called pure dense phase spectrum, which has the lowest root mean sqaure deviation from the spectrum detected in presence of 70% w/v sucrose as shown in the inset on the right (black: difference spectrum, dotted black: 70% sucrose). The A_zz_ is found to be the same in the pure dense phase and in the reference 70% sucrose spectrum. The inset on the left shows the reconstruction methodology. The absolute integral of the residual between the target sucrose spectrum (50, 60, 70% w/v) and the difference spectrum is plotted vs the fraction of the subtracted dilute-phase spectrum. Two minima of the residuals are observed, correlated with the reconstruction with 15% (or 22%) dilute fraction and a 60% (or 70%) sucrose spectrum best representing the pure dense phase.

From the analysis of the series of spectra at different viscosities, it can be recognized that with the increase of viscosity (Fig. 5A), both the positive peak in the low field region and the splitting of the negative central peak become more evident. Notably, the negative central peak splits at sucrose concentrations ≥50% w/v, and while from 50 to 60% there are still some spectral changes, from 60 to 70% the spectral shape does not change despite the increase in viscosity by one order of magnitude, indicating that the motion of the label has reached the rigid limit. We found that the spectrum of spin-labeled crystallin in the dense phase best corresponds to the rigid limit spectrum obtained in 60% w/v sucrose (48.49 mPa·s) from the viscosity ‘ruler’ (Fig 5C). To identify both the residual fraction of dilute phase present in this spectrum and the viscosity which best represented the pure dense phase, we used a reconstruction methodology (see experimental methods) using the dilute-phase spectrum and the viscosity ruler (Fig. 5D). We found that the optimal reconstruction was obtained with both the 60% and 70% sucrose spectra and 15% or 22% residual dilute phase, respectively. Interestingly, the spin-labeled sidechain in the condensed phase experienced the same polarity (encoded in the hyperfine parameter A_zz_) as in the 60-70% w/v sucrose solution.

As the viscosity is linearly correlated with the rotational correlation time (see for example cw EPR spectra obtained with the free spin label MAP in different sucrose solutions in Fig. S3C,D), we can infer that γD-crystallin in the condensed phase at 293 K in presence of 0.6 M TMAO experiences a viscosity close to the rigid limit already reached at 60% sucrose (48.49 mPa·s). Therefore, as lower bound we can estimate that of a 50% w/v sucrose solution (ca. 15 mPa·s), which means that the lower bound for the rotational correlation time is 15 times higher than a very dilute solution in water (i.e. > 190 ns using the value of 12.2 ns determined by fluorescence anisotropy at 293 K). This conclusion was corroborated by the same EPR analysis performed with another protein variant (C111, see Fig. S8), showing an optimal reconstruction with the 60% sucrose spectrum. The 15 times lower bound of retardation is in good agreement with the MD-predicted viscosity increase by more than one order of magnitude.

Finally, we used Double Electron-Electron Resonance (DEER) to investigate the effect of LLPS on the intra-distance between the two spin-labeled labels (S73C-C111) to address the structural integrity of the proteins in condensates in presence of cosolutes and we performed THz spectroscopy to compare the changes in the hydration waters in presence of cosolutes with those previously detected on pure proteins.^53^ Notably, as shown by the MD simulations in the dense phase at protein concentration of 420 mg/ml, the water concentration is ca. 600 mg/ml. The amount of water molecules surrounding each protein is therefore (more than) sufficient to fully hydrate a folded protein. Interestingly, by comparing the intra-protein distance in the dilute phase in presence of deuterated glycerol and 5% w/v PEG, we found that PEG induces a broadening of the distance distribution. This broadening is not due to protein aggregation, rather we suggest it to be a direct effect of the cosolute on the distribution of the spin-labeled rotamers (Fig. S9). In the presence of LLPS we detected a faster phase memory time (Fig. S10) due to higher local spin-labeled protein concentration, a more pronounced background decay in the primary DEER traces (Fig. S9C), but the mean intra-protein distance was maintained (Fig. S9B and Fig. 9C), indicating that the globular fold of crystallin persists in the condensate. The distance distribution is consistent with distance distributions predicted via a rotamer analysis on protein snapshots extracted from the MD simulations in the dilute and condensed phases (Fig. S9 and S11).

The changes in the solvation waters upon formation of LLPS in presence of cosolutes were also found to be consistent with the effects previously described (Fig. S12), which were shown to be in agreement with the solvation information extracted from the same MD simulations used in this study. ^23^

## Conclusions

Accessing the dynamics of proteins undergoing LLPS might be crucial to understand a manifold of biological processes. Here we demonstrated an integrative approach to combine MD simulations, fluorescence and EPR spectroscopy to elucidate the rotational dynamics of the globular protein γD-crystallin. We predicted via MD simulations a rotational correlation time of the protein in aqueous solution of ca 17 ns at 293 K (Table 2), which was validated by fluorescence anisotropy (12.2 ns, Table 2), and a heterogeneous slow-down of the overall tumbling of the crystallin proteins by one to two orders of magnitude for condensed phases of 425 mg/ml, which was corroborated by EPR data.

The continuous wave X-band EPR spectra of nitroxide probes are sensitive reporters of rotational correlation times in a three-order of magnitude range, from ca. 30 ps to ca. 100 nanoseconds (Fig. S3). For globular proteins, the already slow rotational dynamics in the dilute phase decreases the contrast between dilute and condensed phases in terms of EPR spectral effects and therefore prevent a quantitative analysis of the rotational correlation times in terms of spectral deconvolution. However, we demonstrate that it is possible to extract the onset temperature of LLPS of globular proteins such as crystallin in presence of cosolutes by comparing the spectral features of nitroxide-labeled proteins at the same concentrations in the presence and absence of cosolutes. We identified a population of nitroxide probes characterized by slow motion (rotational correlation time > 100 ns), which appears in presence of droplets and is caused by the interactions between the labeled side chain and adjacent proteins. This implies that there is a detectable spectral fingerprint which reports on the rotational dynamics of the spin-labeled side chains in the droplets, characterized by motions approaching the rigid limit of X-band EPR.

Additionally, we used spin-labeled crystallin as viscosity nanoprobe, to correlate the bulk viscosity effects induced by sucrose on a series of spectra to the effects induced by molecular crowding in the droplets. We could show that in the absence of LLPS at concentrations of 1.5 mM the proteins already experience an increased dynamic viscosity of 1.59 mPa·s. We set a threshold for the retardation factor, which is in line with the MD value (Table 2). The nitroxide spectrum in the droplets has clearly reached the rigid limit at X-band frequency, indicating that not only the overall tumbling of the protein, but also the rotational freedom of the side chain has experienced a strong retardation with respect to the dilute phase. The highly restricted motional freedom of the protein’s side chains in a globular protein in the dense phase is a clear effect due to the crowding exerted by the neighbouring proteins in the condensates and might be important in modulating enzymatic activities and interaction propensity of globular proteins in condensate. Interestingly, the small molecular crowder sucrose (molecular weight 342 Da), similar in size to the nitroxide spin labels and the amino acids, was shown to induce spectral effects almost identical to those induced by the protein crowding in the droplets in terms of spectral features and also local polarity (we found the same polarity in high sucrose solutions and in the condensed phase). Therefore, creating a viscosity ‘ruler’ using a spin-labeled protein in presence of molecular solutes such as sucrose can be a useful approach to monitor the microviscosity of the environment in droplets and to identify possible changes due to addition of binding partners in the condensed phase.

## Supporting Information

The supporting information contains: Table S1 with rotational correlation times extracted by MD simulations; Fig. S1 with rotational correlation times versus interaction energy; Fig. S2 with lifetime of fluoresceine-labeled proteins; Fig. S3 with examples of the effects of different rotational correlation times on the EPR spectra; Table S2 with EPR simulation parameters; Fig. S4 with turbidity plots; Fig. S5 with the EPR methods to extract the onset temperature of LLPS; Fig. S6 with the cw X-band EPR spectra; Table S3 with EPR simulation parameters; Fig. S7 with EPR spectra at different protein concentrations; Fig. S8 with the viscosity ruler on the mutant C111; Fig. S9 with the DEER data; Fig. S10 with the phase memory time traces; Fig. S11 with the rotamers calculated on the MD trajectories; Fig. S12 with the THz data.

## Acknowledgement

We would like to thank Sashary Ramos and Martina Havenith from Ruhr University Bochum for the analysis of the samples with THz spectroscopy. This work is supported by the SNF grant number 188817. This project received funding from the European Union’s Horizon 2020 research and innovation programme under the Marie Skłodowska-Curie Grant Agreement No. 801459 -FP-RESOMUS and was funded by the Deutsche Forschungsgemeinschaft (DFG) under Germany’s Excellence Strategy -EXC 2033 -390677874 -RESOLV.

## Abbreviations

LLPS: liquid-liquid phase separation; EPR: electron paramagnetic resonance

## REFERENCES

(1) Zeng, M.; Chen, X.; Guan, D.; Xu, J.; Wu, H.; Tong, P.; Zhang, M. Reconstituted Postsynaptic Density as a Molecular Platform for Understanding Synapse Formation and Plasticity. Cell 2018, 174 (5), 1172–1187.

(2) Kato, M.; Han, T. W.; Xie, S.; Shi, K.; Du, X.; Wu, L. C.; Mirzaei, H.; Goldsmith, E. J.; Longgood, J.; Pei, J.;, et al. Cell-free Formation of RNA Granules: Low Complexity Sequence Domains Form Dynamic Fibers within Hydrogels. Cell 2012, 149 (4), 753–767.

(3) Feric, M.; Vaidya, N.; Harmon, T. S.; Mitrea, D. M.; Zhu, L.; Richardson, T. M.; Kriwacki, R. W.; Pappu, Rohit V.; Brangwynne, Clifford P. Coexisting Liquid Phases Underlie Nucleolar Subcompartments. Cell 2016, 165 (7), 1686–1697.

(4) Riback, J. A.; Katanski, C. D.; Kear-Scott, J. L.; Pilipenko, E. V.; Rojek, A. E.; Sosnick, T. R.; Drummond, D. A. Stress-Triggered Phase Separation Is an Adaptive, Evolutionarily Tuned Response. Cell 2017, 168 (6), 1028–1040.e1019.

(5) Nozawa, R.-S.; Yamamoto, T.; Takahashi, M.; Tachiwana, H.; Maruyama, R.; Hirota, T.; Saitoh, N. Nuclear microenvironment in cancer: Control through liquid-liquid phase separation. Cancer Science 2020, 111 (9), 3155–3163.

(6) Strom, A. R.; Emelyanov, A. V.; Mir, M.; Fyodorov, D. V.; Darzacq, X.; Karpen, G. H. Phase separation drives heterochromatin domain formation. Nature 2017 547:7662 2017, 547 (7662), 241–245.

(7) Zhang, H.; Ji, X.; Li, P.; Liu, C.; Lou, J.; Wang, Z.; Wen, W.; Xiao, Y.; Zhang, M.; Zhu, X.;, et al. Liquid-liquid phase separation in biology: mechanisms, physiological functions and human diseases. Science China Life Sciences 2020, 63 (7), 953–985.

(8) Miesch, J.; Wimbish, R. T.; Velluz, M.-C.; Aumeier, C. Phase separation of +TIP networks regulates microtubule dynamics. Proceedings of the National Academy of Sciences 2023, 120 (35), e2301457120.

(9) Brangwynne, C. P.; Eckmann, C. R.; Courson, D. S.; Rybarska, A.; Hoege, C.; Gharakhani, J.; Jülicher, F.; Hyman, A. A. Germline P granules are liquid droplets that localize by controlled dissolution/condensation. Science (New York, N.Y.) 2009, 324 (5935), 1729–1732.

(10) Jülicher, F.; Weber, C. A. Droplet Physics and Intracellular Phase Separation. Annual Review of Condensed Matter Physics 2024, 15 (Volume 15, 2024), 237–261.

(11) Wakisaka, A.; Ohki, T. Phase separation of water-alcohol binary mixtures induced by the microheterogeneity. Faraday discussions 2005, 129, 231–245.

(12) Bertei, A.; Chueh, C.-C.; Mauri, R. Dynamics of phase separation of sheared binary mixtures after a nonisothermal quenching. Physical Review Fluids 2021, 6 (9), 094302.

(13) Cates, M. E.; Tjhung, E. Theories of binary fluid mixtures: from phase-separation kinetics to active emulsions. Journal of Fluid Mechanics 2018, 836, P1.

(14) Lin, Y.-H.; Forman-Kay, J. D.; Chan, H. S. Theories for Sequence-Dependent Phase Behaviors of Biomolecular Condensates. Biochemistry 2018, 57 (17), 2499–2508.

(15) M. Ruff, K. R. Stefan; Chilkoti, Ashutosh; Pappu, Rohit V. Advances in Understanding Stimulus-Responsive Phase Behavior of Intrinsically Disordered Protein Polymers. Journal of Molecular Biology 2018, 430 (23), 4619–4635.

(16) Brangwynne, Clifford P.; Tompa, P.; Pappu, Rohit V. Polymer physics of intracellular phase transitions. Nature Physics 2015, 11 (11), 899–904.

(17) Broide, M. L.; Tominc, T. M.; Saxowsky, M. D. Using phase transitions to investigate the effect of salts on protein interactions. Physical Review E 1996, 53 (6), 6325–6335.

(18) Braun, M. K.; Sauter, A.; Matsarskaia, O.; Wolf, M.; Roosen-Runge, F.; Sztucki, M.; Roth, R.; Zhang, F.; Schreiber, F. Reentrant Phase Behavior in Protein Solutions Induced by Multivalent Salts: Strong Effect of Anions Cl-Versus NO3. The journal of physical chemistry. B 2018, 122 (50), 11978–11985.

(19) Hansen, J.; Uthayakumar, R.; Pedersen, J. S.; Egelhaaf, S. U.; Platten, F. Interactions in protein solutions close to liquid–liquid phase separation: ethanol reduces attractions via changes of the dielectric solution properties. Physical Chemistry Chemical Physics 2021, 23 (39), 22384–22394.

(20) Chowdhury, A.; Borgia, A.; Ghosh, S.; Sottini, A.; Mitra, S.; Eapen, R. S.; Borgia, M. B.; Yang, T.; Galvanetto, N.; Ivanović, M. T.;, et al. Driving forces of the complex formation between highly charged disordered proteins. Proceedings of the National Academy of Sciences 2023, 120 (41), e2304036120.

21. Hong, J.; Xiong, S. TMAO-Protein Preferential Interaction Profile Determines TMAO’s Conditional In Vivo Compatibility. Biophysical Journal 2016, 111 (9), 1866–1875.

(22) Wu, J.; Zhao, C.; Lin, W.; Hu, R.; Wang, Q.; Chen, H.; Li, L.; Chen, S.; Zheng, J. Binding characteristics between polyethylene glycol (PEG) and proteins in aqueous solution. Journal of Materials Chemistry B 2014, 2 (20), 2983–2992.

(23) T Arakawa; Timasheff, S. N. Mechanism of poly(ethylene glycol) interaction with proteins. Biochemistry 1985, 24 (24), 6756–6762.

(24) Mukherjee, S.; Ramos, S.; Pezzotti, S.; Kalarikkal, A.; Prass, T. M.; Galazzo, L.; Gendreizig, D.; Barbosa, N.; Bordignon, E.; Havenith, M.;, et al. Entropy Tug-of-War Determines Solvent Effects in the Liquid–Liquid Phase Separation of a Globular Protein. The Journal of Physical Chemistry Letters 2024, 15 (15), 4047–4055.

(25) Mukherjee, S.; Schäfer, L. V. Thermodynamic forces from protein and water govern condensate formation of an intrinsically disordered protein domain. Nature Communications 2023, 14 (1), 5892.

(26) Galvanetto, N.; Ivanović, M. T.; Chowdhury, A.; Sottini, A.; Nüesch, M. F.; Nettels, D.; Best, R. B.; Schuler, B. Extreme dynamics in a biomolecular condensate. Nature 2023, 619 (7971), 876–883.

(27) Dindo, M.; Bevilacqua, A.; Laurino, P. Enzymes and Liquid-Liquid Phase Separation: A New Era for the Regulation of Enzymatic Activity. Seibutsu Butsuri 2023, 63 (1), 12–15.

(28) Lyon, A. S.; Peeples, W. B.; Rosen, M. K. A framework for understanding the functions of biomolecular condensates across scales. Nature Reviews Molecular Cell Biology 2020 22:3 2020, 22 (3), 215–235.

(29) Fischer, A. A. M.; Robertson, H. B.; Kong, D.; Grimm, M. M.; Grether, J.; Groth, J.; Baltes, C.; Fliegauf, M.; Lautenschläger, F.; Grimbacher, B.;, et al. Engineering Material Properties of Transcription Factor Condensates to Control Gene Expression in Mammalian Cells and Mice. Small 2024, 2311834.

(30) Peran, I. M., Tanja Molecular structure in biomolecular condensates. Current opinion in structural biology 2020, 60, 17–26.

(31) Vendra, V. P. R.; Khan, I.; Chandani, S.; Muniyandi, A.; Balasubramanian, D. Gamma crystallins of the human eye lens. Biochimica et Biophysica Acta (BBA) -General Subjects 2016, 1860 (1), 333–343.

(32) Basak, A.; Bateman, O.; Slingsby, C.; Pande, A.; Asherie, N.; Ogun, O.; Benedek, G. B.; Pande, J. High-resolution X-ray Crystal Structures of Human γD Crystallin (1.25Å) and the R58H Mutant (1.15Å) Associated with Aculeiform Cataract. Journal of Molecular Biology 2003, 328 (5), 1137–1147.

(33) Cinar, S.; Cinar, H.; Chan, H. S.; Winter, R. Pressure-Sensitive and Osmolyte-Modulated Liquid-Liquid Phase Separation of Eye-Lens γ-Crystallins. Journal of the American Chemical Society 2019, 141 (18), 7347–7354.

(34) Cinar, H.; Winter, R. The effects of cosolutes and crowding on the kinetics of protein condensate formation based on liquid–liquid phase separation: a pressure-jump relaxation study. Scientific Reports 2020, 10 , 17245.

(35) Wang, Y.; Lomakin, A.; McManus, J. J.; Ogun, O.; Benedek, G. B. Phase behavior of mixtures of human lens proteins Gamma D and Beta B1. Proceedings of the National Academy of Sciences 2010, 107 (30), 13282–13287.

(36) Zhao, H.; Magone, M. T.; Schuck, P. The role of macromolecular crowding in the evolution of lens crystallins with high molecular refractive index. Physical Biology 2011, 8 (4), 046004.

(37) Uhlhorn, S. R.; Borja, D.; Manns, F.; Parel, J.-M. Refractive index measurement of the isolated crystalline lens using optical coherence tomography. Vision Research 2008, 48 (27), 2732–2738.

(38) Chen, J.; Callis, P. R.; King, J. Mechanism of the Very Efficient Quenching of Tryptophan Fluorescence in Human γD-and γS-Crystallins: The γ-Crystallin Fold May Have Evolved To Protect Tryptophan Residues from Ultraviolet Photodamage. Biochemistry 2009, 48 (17), 3708–3716.

(39) Hemmingsen, J. M.; Gernert, K. M.; Richardson, J. S.; Richardson, D. C. The tyrosine corner: A feature of most greek key β-barrel proteins. Protein Science 1994, 3 (11), 1927– 1937.

(40) Harding, J. J.; Dilley, K. J. Structural proteins of the mammalian lens: A review with emphasis on changes in development, aging and cataract. Experimental Eye Research 1976, 22 (1), 1–73.

(41) Zhao, H.; Brown, P. H.; Schuck, P. On the Distribution of Protein Refractive Index Increments. Biophysical Journal 2011, 100 (9), 2309–2317.

(42) Truscott, R. J. W. Age-related nuclear cataract—oxidation is the key. Experimental Eye Research 2005, 80 (5), 709–725.

(43) Moreau, K. L.; King, J. A. Protein misfolding and aggregation in cataract disease and prospects for prevention. Trends in Molecular Medicine 2012, 18 (5), 273–282.

(44) Petitt, P.; Forciniti, D. Cold cataracts: a naturally occurring aqueous two-phase system. Journal of Chromatography B: Biomedical Sciences and Applications 2000, 743 (1-2), 431–441.

(45) Seal, M.; Weil-Ktorza, O.; Despotović, D.; Tawfik, D. S.; Levy, Y.; Metanis, N.; Longo, L. M.; Goldfarb, D. Peptide-RNA Coacervates as a Cradle for the Evolution of Folded Domains. Journal of the American Chemical Society 2022, 144 (31), 14150–14160.

(46) Seal, M.; Jash, C.; Jacob, R. S.; Feintuch, A.; Harel, Y. S.; Albeck, S.; Unger, T.; Goldfarb, D. Evolution of CPEB4 Dynamics Across its Liquid–Liquid Phase Separation Transition. The Journal of Physical Chemistry B 2021, 125 (47), 12947–12957.

(47) Bramham, J. E.; Golovanov, A. P. Temporal and spatial characterisation of protein liquid-liquid phase separation using NMR spectroscopy. Nature Communications 2022, 13 (1), 1767.

(48) Esteban-Hofer, L.; Emmanouilidis, L.; Yulikov, M.; Allain, F. H.-T.; Jeschke, G. Ensemble structure of the N-terminal domain (1–267) of FUS in a biomolecular condensate. Biophysical Journal 2024, 123 (5), 538–554.

(49) Ritsch, I.; Lehmann, E.; Emmanouilidis, L.; Yulikov, M.; Allain, F.; Jeschke, G. Phase Separation of Heterogeneous Nuclear Ribonucleoprotein A1 upon Specific RNA-Binding Observed by Magnetic Resonance. Angewandte Chemie International Edition 2022, 61 (40), e202204311.

(50) Robustelli, P.; Piana, S.; Shaw, D. E. Developing a molecular dynamics force field for both folded and disordered protein states. Proceedings of the National Academy of Sciences 2018, 115 (21), 4758–4766.

(51) Hoover, W. G. Canonical dynamics: Equilibrium phase-space distributions. Physical Review A 1985, 31 (3), 1695–1697.

(52) Parrinello, M.; Rahman, A. Polymorphic transitions in single crystals: A new molecular dynamics method. Journal of Applied Physics 1981, 52 (12), 7182–7190.

(53) Darden, T.; York, D.; Pedersen, L. Particle mesh Ewald: An N⋅log(N) method for Ewald sums in large systems. The Journal of Chemical Physics 1993, 98 (12), 10089–10092.

(54) Hess, B.; Bekker, H.; Berendsen, H. J. C.; Fraaije, J. G. E. M. LINCS: A linear constraint solver for molecular simulations. Journal of Computational Chemistry 1997, 18 (12), 1463–1472.

(55) Miyamoto, S.; Kollman, P. A. Settle: An analytical version of the SHAKE and RATTLE algorithm for rigid water models. Journal of Computational Chemistry 1992, 13 (8), 952–962.

(56) Chen, P.-c.; Hologne, M.; Walker, O. Computing the Rotational Diffusion of Biomolecules via Molecular Dynamics Simulation and Quaternion Orientations. The Journal of Physical Chemistry B 2017, 121 (8), 1812–1823.

(57) Hoffmann, F.; Xue, M.; Schäfer, L. V.; Mulder, F. A. A. Narrowing the gap between experimental and computational determination of methyl group dynamics in proteins. Physical Chemistry Chemical Physics 2018, 20 (38), 24577–24590.

(58) Linke, M.; Köfinger, J.; Hummer, G. Fully Anisotropic Rotational Diffusion Tensor from Molecular Dynamics Simulations. The Journal of Physical Chemistry B 2018, 122 (21), 5630–5639.

(59) Tschaggelar, R.; Kasumaj, B.; Santangelo, M. G.; Forrer, J.; Leger, P.; Dube, H.; Diederich, F.; Harmer, J.; Schuhmann, R.; García-Rubio, I.;, et al. Cryogenic 35 GHz pulse ENDOR probehead accommodating large sample sizes: Performance and applications. Journal of Magnetic Resonance 2009, 200 (1), 81–87.

(60) Pannier, M.; Veit, S.; Godt, A.; Jeschke, G.; Spiess, H. W. Dead-time free measurement of dipole-dipole interactions between electron spins. Journal of magnetic resonance (San Diego, Calif. : 1997) 2000, 142 (2), 331–340.

(61) E. Tait, C.; Stefan Stoll. Coherent pump pulses in Double Electron Electron Resonance spectroscopy. Physical Chemistry Chemical Physics 2016, 18 (27), 18470–18485.

(62) Jeschke, G.; Chechik, V.; Ionita, P.; Godt, A.; Zimmermann, H.; Banham, J.; Timmel, C. R.; Hilger, D.; Jung, H. DeerAnalysis2006—a comprehensive software package for analyzing pulsed ELDOR data. Applied Magnetic Resonance 2006, 30 (3), 473–498.

(63) Walfort, B.; Gartmann, N. A., Jafar; Rosspeintner, Arnulf; Hagemann, Hans. Effect of excitation wavelength (blue vs near UV) and dopant concentrations on afterglow and fast decay of persistent phosphor SrAl2O4:Eu2+,Dy3+. Journal of Rare Earths 2022, 40 (7), 1022–1028.

(64) Wahl, P. Analysis of fluorescence anisotropy decays by a least square method. Biophysical chemistry 1979, 10 (1), 91–104.

(65) Dar, F.; Cohen, S. R.; Mitrea, D. M.; Phillips, A. H.; Nagy, G.; Leite, W. C.; Stanley, C. B.; Choi, J. M.; Kriwacki, R. W.; Pappu, R. V. Biomolecular condensates form spatially inhomogeneous network fluids. Nat Commun 2024, 15 (1), 3413.

(66) Chen, Y.; Zhao, H.; Schuck, P.; Wistow, G. Solution properties of γ-crystallins: Compact structure and low frictional ratio are conserved properties of diverse γ-crystallins. Protein Science 2014, 23 (1), 76–87.

(67) Fischer, H.; Polikarpov, I.; Craievich, A. F. Average protein density is a molecular-weight-dependent function. Protein Science : A Publication of the Protein Society 2004, 13 (10), 2825–2828.

(68) Stoll, S.; Schweiger, A. EasySpin, a comprehensive software package for spectral simulation and analysis in EPR. Journal of Magnetic Resonance 2006, 178 (1), 42–55.

(69) Bordignon, E. EPR Spectroscopy of Nitroxide Spin Probes. In EPR Spectroscopy: Fundamentals and Methods, Goldfarb, D. Ed.; Wiley, 2018; pp 277–301.

(70) Hubbell, W. L.; Gross, A.; Langen, R.; Lietzow, M. A. Recent advances in site-directed spin labeling of proteins. Current Opinion in Structural Biology 1998, 8 (5), 649–656.

(71) Hwang, J. S.; Mason, R. P.; Hwang, L. P.; Freed, J. H. Electron spin resonance studies of anisotropic rotational reorientation and slow tumbling in liquid and frozen media. III. Perdeuterated 2,2,6,6-tetramethyl-4-piperidone N-oxide and an analysis of fluctuating torques. J. Phys. Chem. 1975, 79 (5), 489–511.

