## Supporting Information for "A Combined Approach to Extract Rotational Dynamics of Globular Proteins undergoing Liquid-Liquid Phase Separation"

**Table S1. Rotational correlation times and diffusion coefficients of γD-crystallin molecules from MD simulations of the dilute and condensate systems.**

| Protein | Rotational correlation time $\tau_c$ [ns] | Diffusion coefficient $D_{iso}$ [$10^5$ $s^{-1}$] |
|---|---|---|
| Protein in dilute system | 30 | 56.0 |
| Condensate system, Protein 1 | 415 | 4.01 |
| Condensate system, Protein 2 | 1485 | 1.12 |
| Condensate system, Protein 3 | 1530 | 1.09 |
| Condensate system, Protein 4 | 3720 | 0.45 |
| Condensate system, Protein 5 | 4000 | 0.42 |
| Condensate system, Protein 6 | 5545 | 0.30 |
| Condensate system, Protein 7 | 6455 | 0.26 |
| Condensate system, Protein 8 | 6575 | 0.25 |

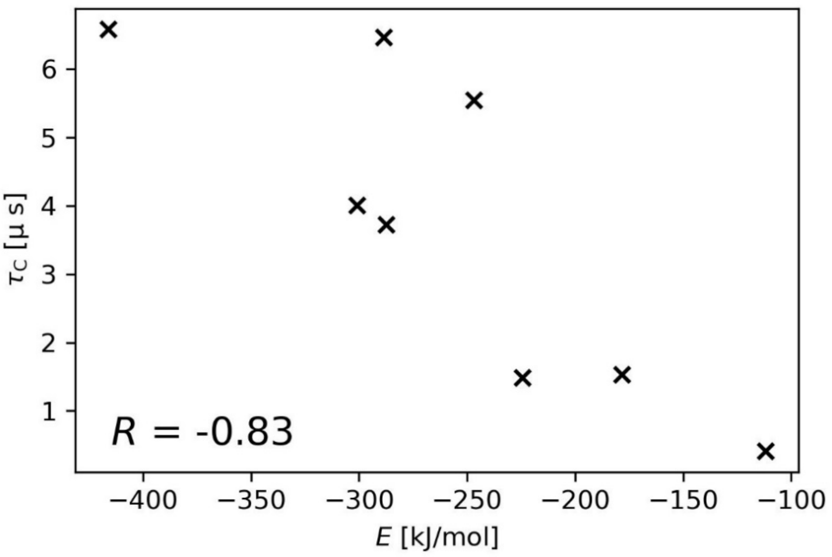


**Figure S1. Isotropic rotational correlation times $\tau_c$ of each protein in the condensate MD system plotted against the average interaction energy $E$ of each protein with all other proteins within 1.5 nm.** The Pearson correlation coefficient $R$ included in the plot shows that $\tau_c$ and $E$ are anticorrelated.

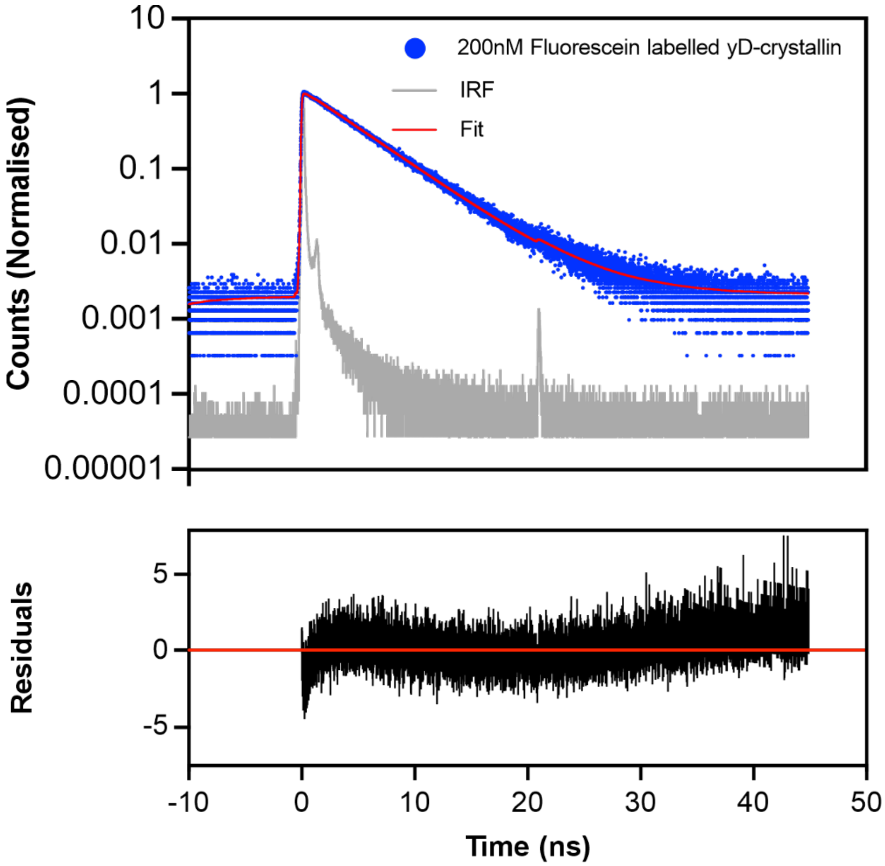


**Figure S2. Lifetime of fluoresceine-labeled protein in aqueous buffer.** Time correlated single photon counting measurement of fluorescein labeled γD-crystallin (200 nM). The histogram of photon counts of the of the fluorescein is shown in blue. The data were fitted with a monoexponential decay function to obtain a fluorescence lifetime of (4.18 ± 0.02) ns. The instrument response function (IRF) is shown in grey. The residuals are shown in black in the bottom plot.

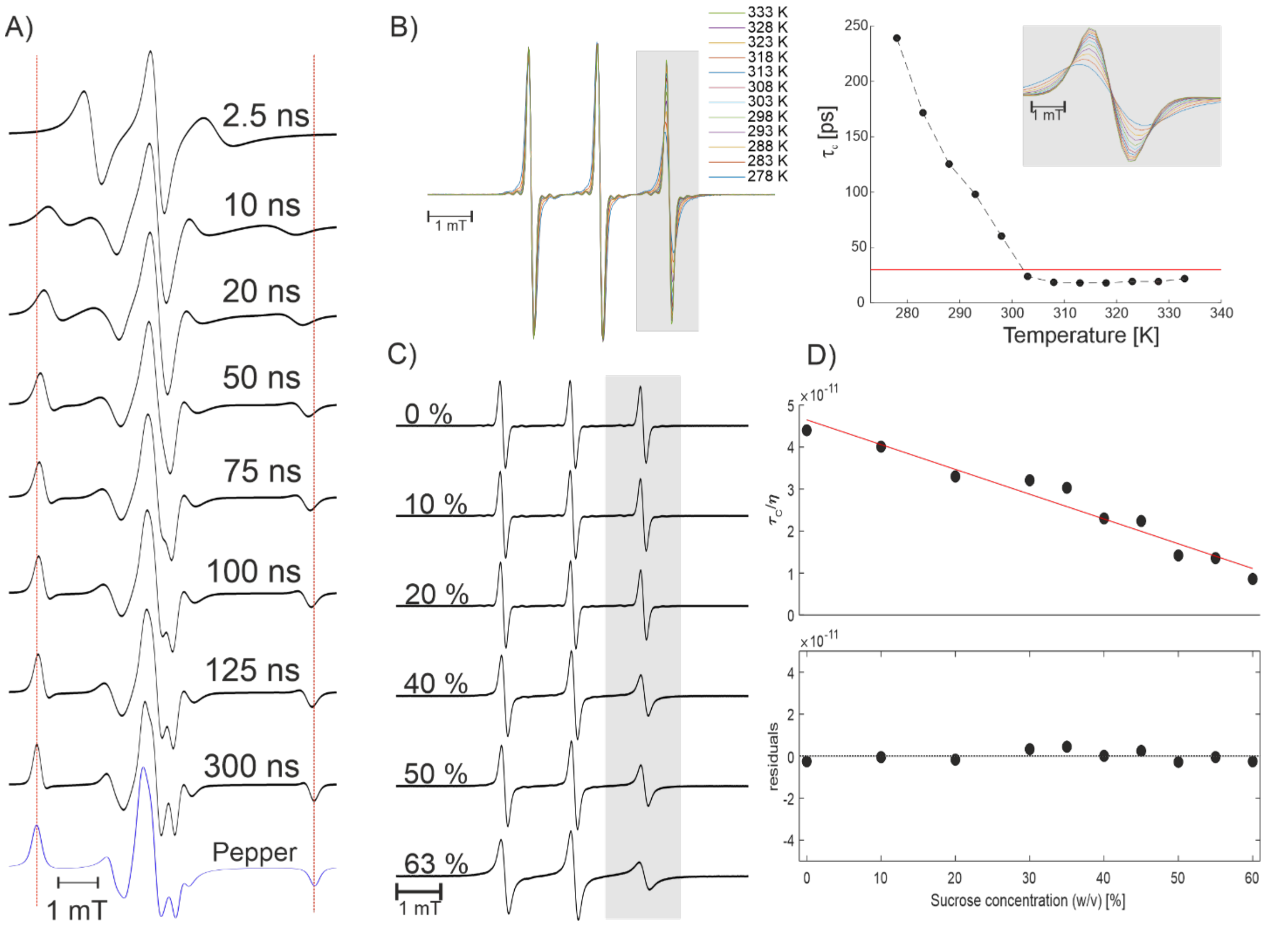


**Figure S3. Influence of the rotational correlation time on the cw EPR X-band spectra.** A) Simulation of nitroxide cw spectra at X-band at different rotational correlation times (EasySpin function chili, isotropic motion[1]) indicating the most sensitive range of rotational dynamics between 2.5 ns and 300 ns with an additional simulation of the rigid limit spectrum (EasySpin function: pepper). Simulation details are shown in table S3. B) Intensity normalized cw EPR spectra of the MAP label in presence of 40 % sucrose at different temperatures. The grey shaded area highlights the part of the spectrum most affected by temperature changes(left). Rotational correlation times plotted against temperature. The high field peak of the spectra is shown in the inlet. The red line indicates a rotational correlation time of 30 ps, the limit of resolution for dynamics at X-band frequency (right). C) The viscosity of the solvent affects the rotational correlation time in a linear fashion. cw EPR spectra of the maleimido-3-proxyl label in presence of different concentrations of sucrose at 293 K. The grey shaded area highlights the most affected part of the spectrum. D) (Top) Rotational correlation times extracted with esfit (EasySpin) divided by the viscosity at the given sucrose concentration, plotted against the content of sucrose. The curve was fitted by linear regression. (Bottom) Residuals analysis of the beforementioned plot shows a random distribution around zero indicating an adequate fit.

**Table S2: Simulation details for the spectra displayed in Fig. S3**

| | *chili* | *pepper* |
|---|---|---|
| $g_x$ | 2.008 | 2.008 |
| $g_y$ | 2.006 | 2.006 |
| $g_z$ | 2.003 | 2.003 |
| $A_{x,y}$ [MHz] | 20 | 20 |
| $A_z$ [MHz] | 95.3 | 95 |
| $lwpp_1$ [mT] | 0.11 | 0.15 |
| $lwpp_2$ [mT] | 0.01 | 0.1 |
| $\tau_c$ [ns] | see Fig. S3 | - |

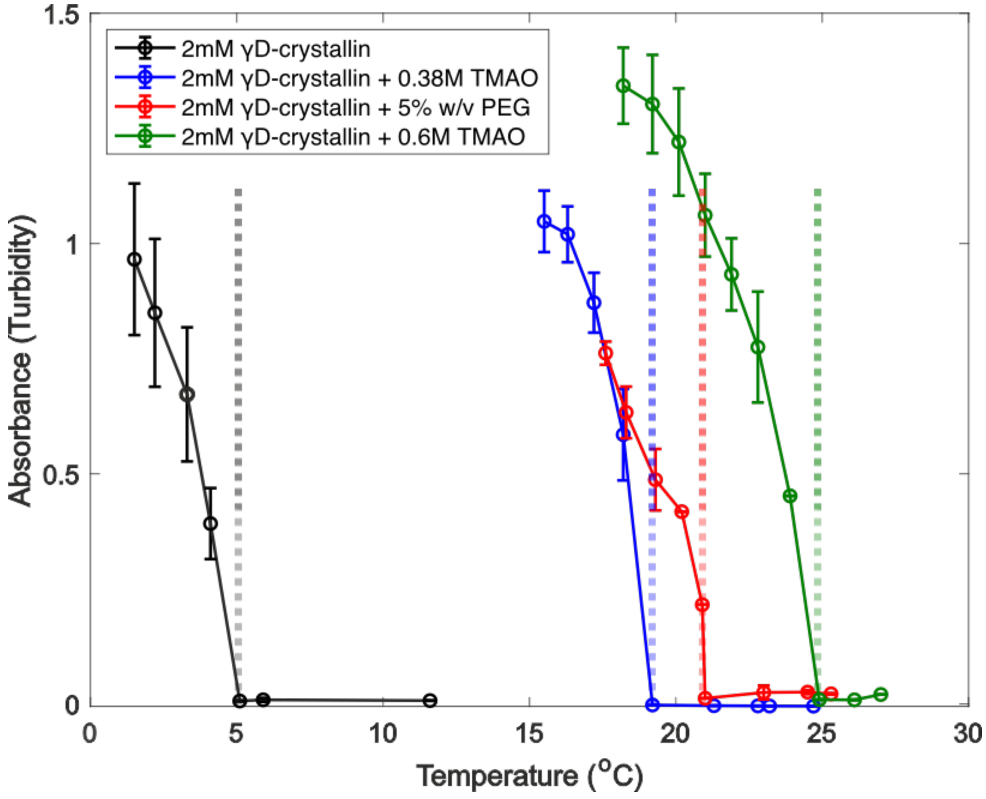


**Figure S4. LLPS onset temperatures by turbidity assays.** The absorbance at 400 nm for samples containing 2mM γD-crystallin was recorded at different temperatures and in the presence of the cosolute trimethylamine-N-oxide (TMAO) and crowding agent polyethylene glycol (PEG). The coloured dashed lines indicate the last measured temperature before the change in the absorbance (turbidity) is observed (LLPS onset temperature). The onset temperature is 5°C for 2 mM yD-crystallin in buffer (20 mM TRIS, 150 mM NaCl, pH = 7.5) and it increases to19, 22.5 and 25°C upon addition of 0.38 M TMAO, 5% w/v PEG and 0.6 M TMAO, respectively

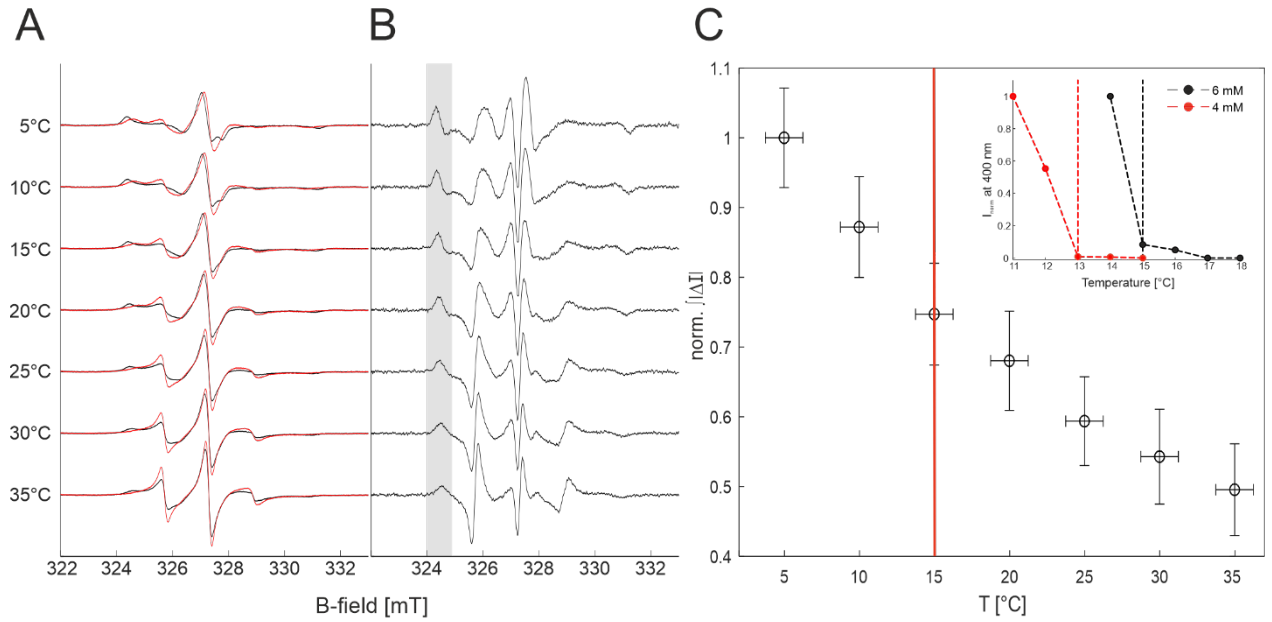


**Figure S5. Determination of the LLPS onset temperature of 5.7 mM γD-crystallin with 50 µM spin-labeled γD-crystallin (C111S-S73C).** A) Overlay of 50 µM spin-labeled γD-crystallin (red) and 50 µM spin-labeled γD-crystallin with 5.7 mM unlabeled γD-crystallin (black). B) Difference spectra (black minus red in panel A) at the respective temperatures. The grey shaded area highlights the low field area of the spectrum, most sensitive to changes of mobility C) Normalized absolute integral extracted from the differential spectra shown in B). The red line indicates the onset temperature of approximately 15 °C under these conditions. The contrast between the reference and the sample is not adequate to extract the onset temperature via cw EPR. The Inlet shows the onset temperature of pure protein at 4 mM (red) and 6 mM (black) concentration determined by turbidity assay.

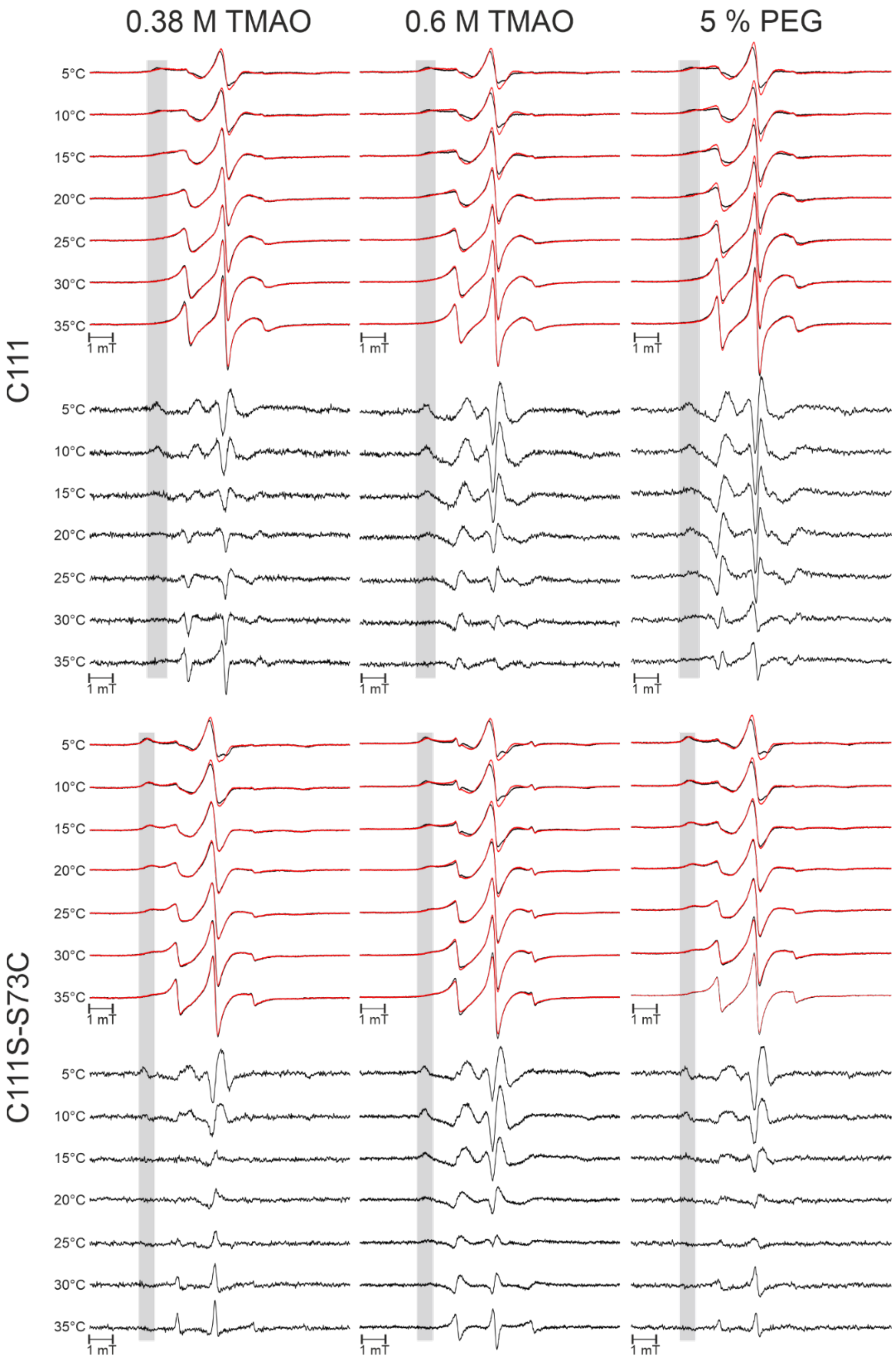


**Figure S6**. **X-band cw EPR spectra recorded at different temperatures in presence and absence of cosolutes.** Cw EPR spectra of samples containing 2 mM γD-crystallin and 50 µM spin-labeled protein at different temperatures in absence (reference spectra, red) and in presence of cosolute (black). The grey shaded area denotes the low field region of the spectra in which an "immobile" positive component appears upon LLPS formation. Difference spectra are shown below and were obtained subtracting the reference spectra from the spectra with added co-solute at the corresponding temperature. The grey shaded area highlights the "immobile" positive spectral feature that was integrated to create Figure 4.

**Table S3: Details for the simulations of the dilute phase spectrum in Fig. 5**

| | Component 1 | Component 2 |
|---|---|---|
| $g_x$ | 2.0087 | 2.0087 |
| $g_y$ | 2.0063 | 2.0063 |
| $g_z$ | 2.0023 | 2.0023 |
| $A_{x,y}$ [MHz] | 20 | 20 |
| $A_z$ [MHz] | 104.9 | 104.9 |
| $lwpp_1$ [mT] | 0.11 | 0.11 |
| $lwpp_2$ [mT] | 0.01 | 0.01 |
| $\tau_c$ [ns] | 2.51 | 8.82 |
| Weight | 0.295 | 0.705 |

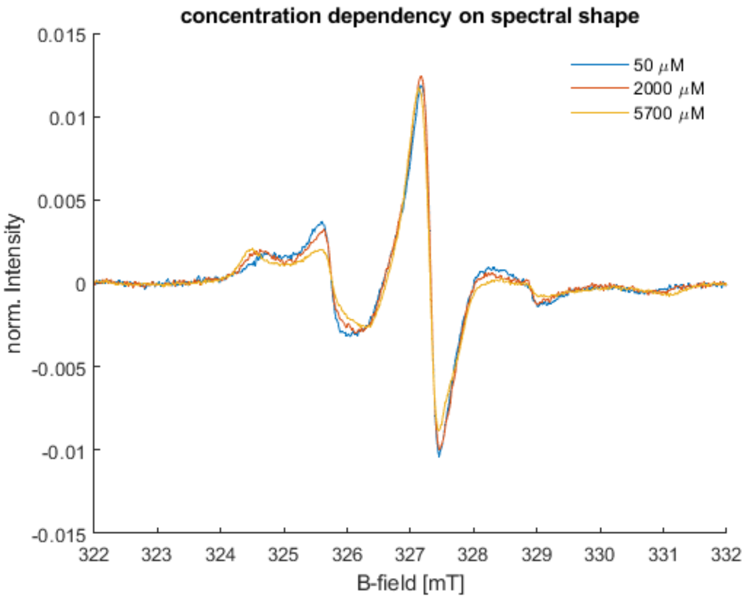


**Figure S7**. Comparison of the spectral shape of 50 µM spin-labeled γD-crystallin (C111S-S73C) in presence of varying concentrations of unlabeled wt γD-crystallin at 293 K. The spectra are normalized to the second integral (see M&M) With increasing concentration the spectrum shows features of a more immobile protein due to crowding effects.

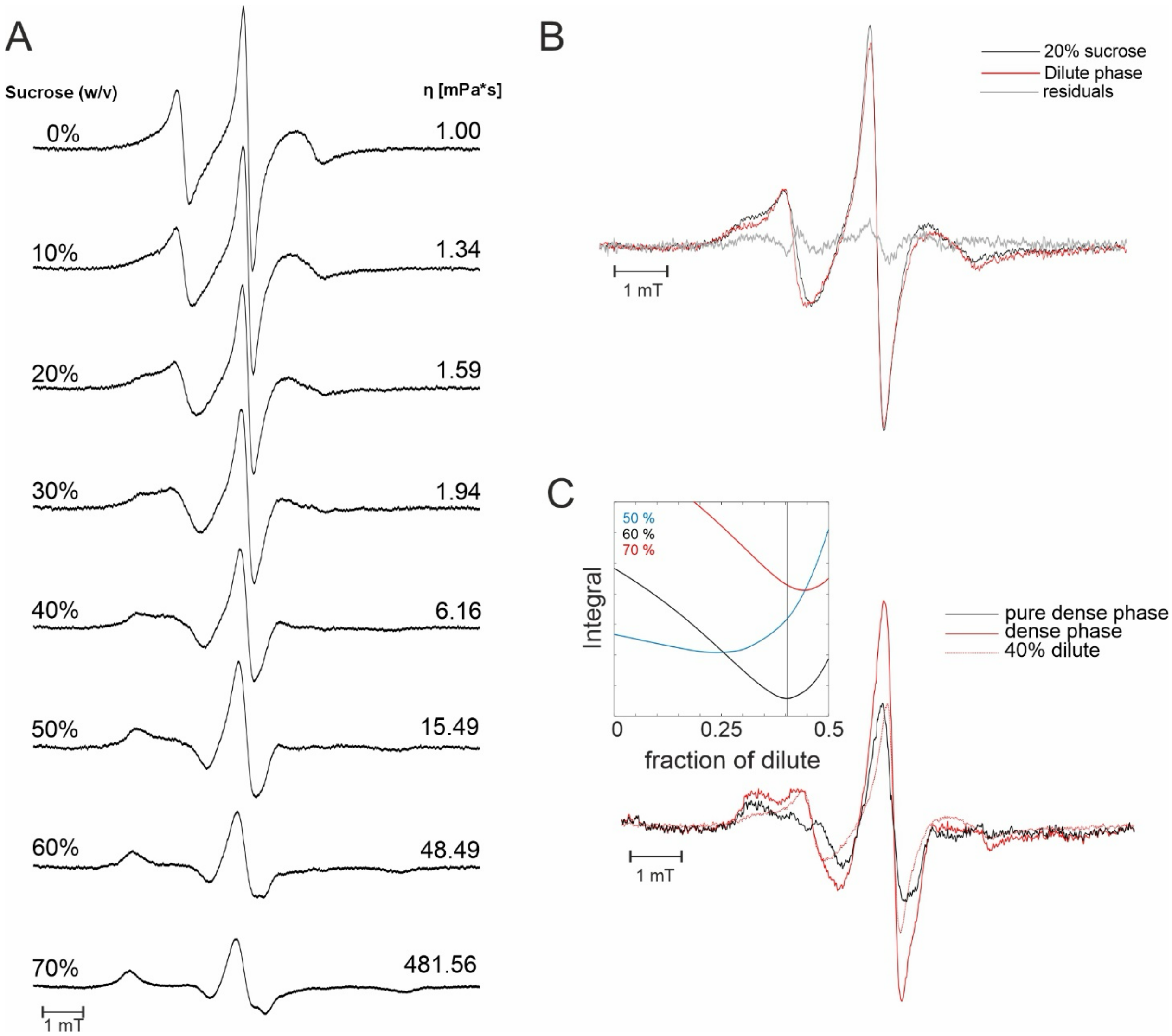


**Figure S8**. Viscosity 'ruler' and spectral reconstruction of spin-labeled γD-crystallin C111. A) Cw EPR spectra of γD-crystallin (C111) recorded at 293 K in aqueous buffer with increasing sucrose concentrations. The corresponding tabulated dynamic viscosity of sucrose solutions at 293 K is indicated on the right. Similarly to the sucrose series obtained with the C111S-S73 (Fig. 5), the rigid limit is reached above 60% sucrose concentration. B) The cw EPR spectrum of γD-crystallin (C111) in the dilute phase (red) best corresponds to the spectrum of the same mutant in 20% sucrose solutions (1.59 mPas viscosity, black, from panel A), confirming the results obtained with the mutant C111S-S73C (Fig. 5). The residual is shown in grey. C) From the dense-phase spectrum (red) 40% of the dilute phase spectrum was subtracted (red dotted, as in panel B). The obtained difference spectrum (black) yields the so-called pure dense phase spectrum, which has the lowest root mean sqaure deviation from the spectrum detected in presence of 60% w/v sucrose as shown in the inset on the left. The inset on the left shows the reconstruction methodology. The absolute integral of the residual between the target sucrose spectrum (50, 60, 70% w/v) and the difference spectrum is plotted vs the subtracted fraction of dilute-phase spectrum. Two minima of the residuals are observed with 60 and 70% sucrose as in Fig. 5D. However, the deviations are the smallest for the reconstruction with 40% dilute fraction and a 60% sucrose spectrum representing best the pure dense phase. This fully corroborates the results presented in Fig. 5.

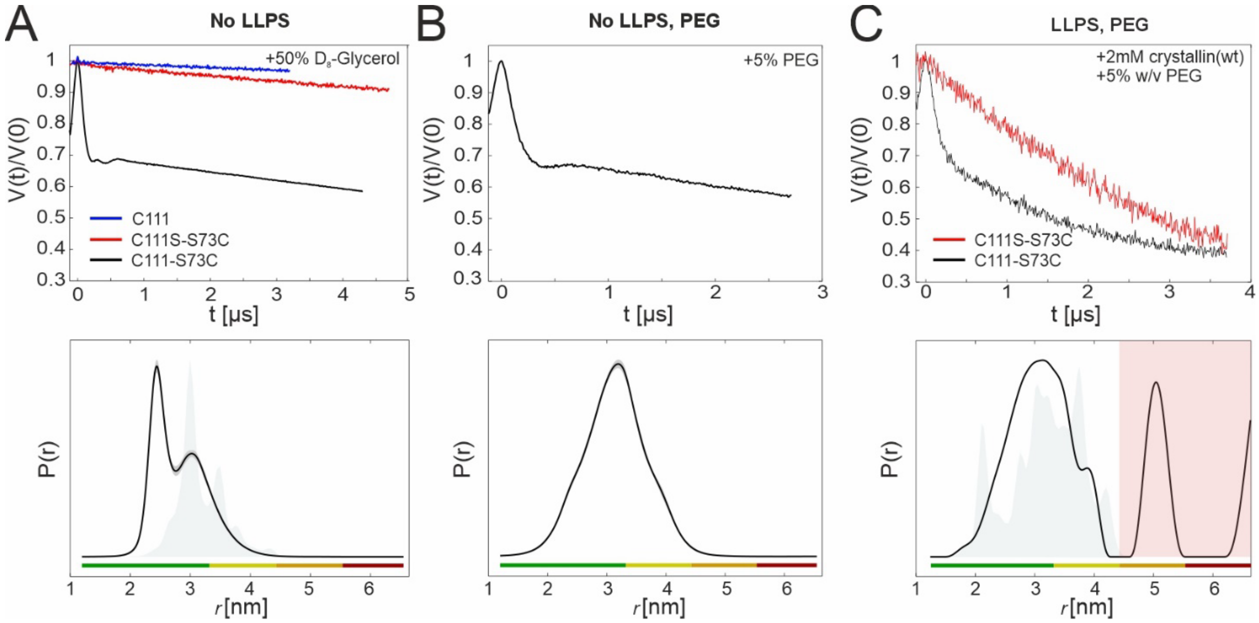


**Figure S9. DEER of spin-labeled γD-crystallin variants under different conditions**. A) Primary DEER traces (upper panel) and distance distributions analyzed with DEERNet[2] within DeerAnalysis[3]of 50 µM spin-labeled crystallin samples in absence of LLPS. The protein with two accessible cysteines (C111-S73C) shows a dipolar oscillation, and a bimodal distance distribution. The shaded grey area is the simulated distance distribution (calculated with MMM 2018.2[3,4]) on the MD trajectory of the monomeric protein structure (rotamers presented in Fig. S11). The DEER traces obtained on the C111 and C111S-S73C protein variants show no dipolar oscillations, indicating that the protein is monomeric and no aggregates are formed. B) Primary DEER trace (upper panel) and distance distribution of 50 µM spin-labeled crystallin (C111-S73C) in presence of 5% w/v PEG (absence of LLPS) analyzed with DEERNet within DeerAnalysis. A broader distance distribution centered at around 3.2 nm is detected in the presence of PEG. C) Primary DEER traces (upper panel) and distance distribution of 50 µM spin-labeled crystallin (C111-S73C, black) analyzed with Tikhonov regularization in DeerAnalysis in presence of 2 mM wild type unlabeled crystallin and 5% w/v PEG (presence of LLPS). The pronounced background decay indicates a higher protein concentration in the presence of LLPS. The distance distribution is similar to that obtained on the monomeric protein solution in the presence of PEG (shown in B). The shaded grey area is the simulated distance distribution (calculated with MMM 2018.2[4,5]) on the MD trajectory of the proteins in the dense phase (425 mg/ml, see Fig. S11). As a control, the DEER trace of the spin-labeled C111S-S73C variant shows as expected a steeper background decay indicating a higher protein concentration in the presence of LLPS, but no signs of aggregates are detected.

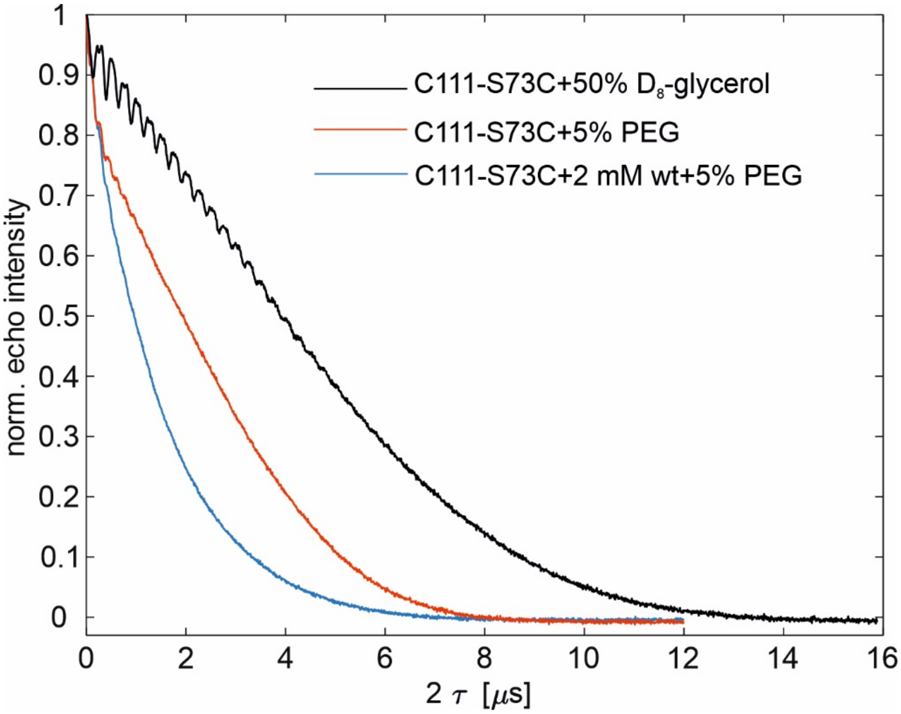


**Figure S10. Phase memory time of the C111-S73C γD-crystallin under different experimental conditions.** The phase memory time traces were recorded at Q band, at 50 K using a 36(π/2)-72(π) ns Hahn-echo sequence with Gaussian pulses (same length and shape used for the DEER traces) with a starting interpulse delay of 160 ns incremented by τ. The graph shows the echo intensity vs 2τ. In black the 50 μM monomeric protein in presence of 50% v/v deuterated glycerol, in red the 50 μM monomeric protein in presence of 5% w/v PEG (no LLPS) and in blue the 50 μM monomeric protein in presence of 2 mM wild type protein and 5% w/v PEG (LLPS). The formation of the dense phase decreases the phase memory time, as expected due to the higher local spin concentration.

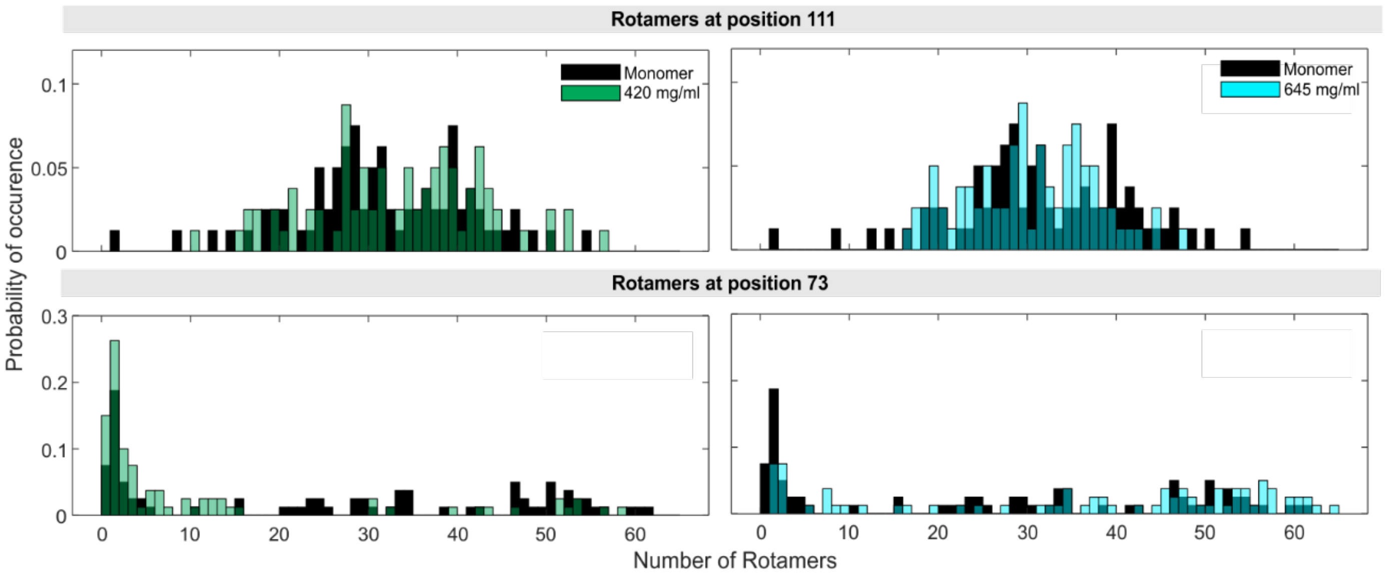


**Figure S11. Analysis of rotamer populations of MAP labeled γD-crystallin in dilute and dense phase from MD simulations used to predict the distance distributions.** (Top panels) Comparison of the rotamer distributions of MAP calculated with MMM 2018.2[4,5] attached to position C111 in γD-crystallin for two different protein densities, 420 mg/ml (green) and 645 mg/ml (cyan) compared to the monomer (black). (Bottom panels) Analysis of the rotamer distribution on position S73C. While the distribution of the rotamers at position C111 does not significantly change between different protein densities, the distribution of the rotamers at position S73C at a density of 645 mg/ml shifts to populations with a higher number of rotamers.

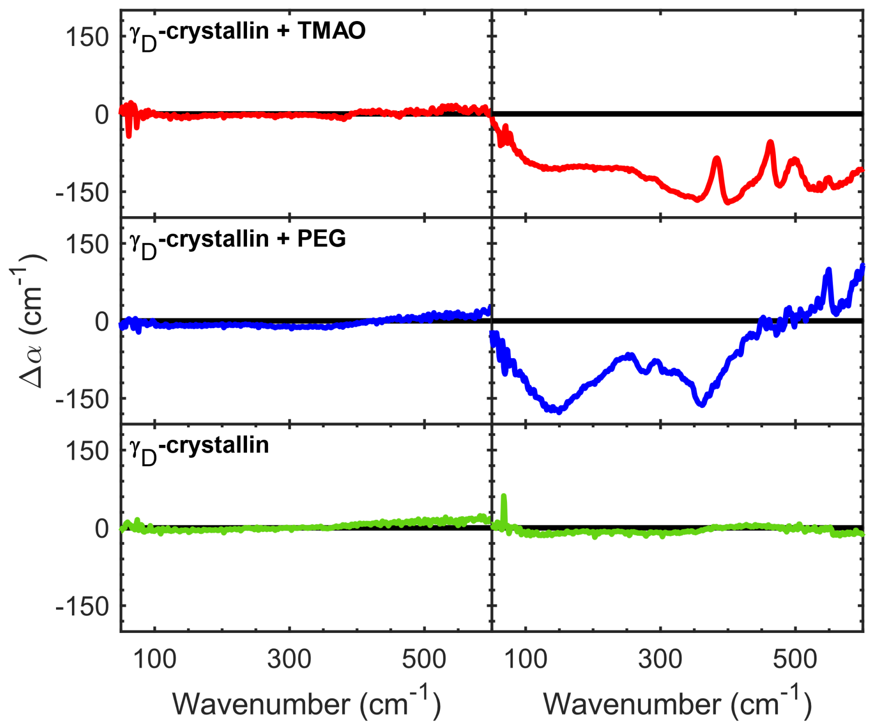


**Figure S12**. **THz spectra of γD-crystallin in presence and absence of cosolutes.** Difference spectra generated by the subtraction of the absorption coefficient of buffer or buffer and crowder solution from that of 2 mM γD-crystallin or 2 γD-mM crystallin with cosolute, $\Delta\alpha_{crystallin} = \alpha_{crystallin}(\nu) - \alpha_{buffer}(\nu)$, at two experimental temperatures 30°C (left) and 15°C (right). γD-crystallin with 0.6 M TMAO (top), with 5% (w/v) PEG, and with no additive (bottom) are shown. No LLPS spectral signatures are observed in crystallin in aqueous solution at either of the probed temperatures. At 30°C, no LLPS signatures are also observed in the spectra with cosolutes (PEG and TMAO) but the fingerprint changes already assigned to the formation of LLPS are evident at 15°C in presence of cosolutes (in agreement with the onset temperature of LLPS of about 20-25°C with 5% PEG and TMAO 0.6 M, see Figure S4). Sharp vibrational signatures from TMAO present in the condensed phase in the top spectra limit the ability to fully disentangle the contributions from the water.

### References


(1) Stoll, S.; Schweiger, A. EasySpin, a comprehensive software package for spectral simulation and analysis in EPR. *J. Magn. Reson.* **2006** 178(1), 42-55
(2) Worswick, S.G. et al. Deep neural network processing of DEER data. *Sci. Adv.* **2018**, 4, eaat521
(3) Jeschke, G.; Chechik, V.; Ionita, P.; Godt, A.; Zimmermann, H.; Banham, J.; Timmel, C. R.; Hilger, D.; Jung, H. DeerAnalysis2006—a comprehensive software package for analyzing pulsed ELDOR data. *Applied Magnetic Resonance* **2006**, *30* (3), 473-498
(4) Y. Polyhach, Y.; Bordignon, E.; Jeschke, G. Rotamer libraries of spin-labelled cysteins for protein studies. *PCCP* **2011** 13(6) 2356-2366
(5) Jeschke, G. MMM: A toolbox for integrative structure modeling. *Protein Sci.* **2018**, **27**, 76-85
